# Precision environmental health monitoring by longitudinal exposome and multi-omics profiling

**DOI:** 10.1101/2021.05.05.442855

**Authors:** Peng Gao, Xiaotao Shen, Xinyue Zhang, Chao Jiang, Sai Zhang, Xin Zhou, Sophia Miryam Schüssler-Fiorenza Rose, Michael Snyder

## Abstract

Conventional environmental health studies primarily focused on limited environmental stressors at the population level, which lacks the power to dissect the complexity and heterogeneity of individualized environmental exposures. Here we integrated deep-profiled longitudinal personal exposome and internal multi-omics to systematically investigate how the exposome shapes an individual’s phenome. We annotated thousands of chemical and biological components in the personal exposome cloud and found they were significantly correlated with thousands of internal biomolecules, which was further cross validated using corresponding clinical data. In particular, our results showed that agrochemicals (e.g., carcinogenic pesticides, fungicides, and herbicides) and fungi predominated in the highly diverse and dynamic personal exposome, and the biomolecules and pathways related to the individual’s immune system, kidneys, and liver were highly correlated with the personal external exposome. Overall, our findings demonstrate dynamic interactions between the personal exposome and internal multi-omics and provide important insights into the impact of the environmental exposome on precision health.

## Introduction

Human health is shaped by the personal genome, microbiome, and exposome [1]. Extensive studies have been conducted on the genome and microbiome, however, the human exposome is rarely investigated, especially at the individual level. Exposomics research aims to characterize all physical, chemical, and biological components collectively in the human external and internal environment. The external environment consists of all potential exposures from the near-field to the far-field sources of exogenous chemical, biological, and physical exposures [2–5]. The internal environment includes but is not limited to dietary components [6–8], xenobiotics and their biotransformation products, foreign DNA/RNA, and bioactive molecules accumulated from exogenous sources [9].

Conventional environmental health risk assessments rely on environmental epidemiology within a predefined, usually large, geographical region. However, recent studies revealed that personal exposome profiles are highly dynamic and spatiotemporally different among individuals who live in the same geographical area. For instance, studies have shown that individuals are exposed to significantly different chemical and biological stressors during the same period even if they are in the same general geographical region, such as the San Francisco Bay Area or London [10, 11]. Another limitation is that previous studies usually targeted a single group of stressors, which failed to provide a holistic picture of the exposome cloud and their interactions [12]. Moreover, stressor-induced physiological responses varied significantly among different individuals [11]. Therefore, there is a critical need to monitor exposures at the individual level and systematically integrate them with respective internal multi-omics profiles to fully characterize each individual’s personal responses to environmental exposures.

Multi-omics analyses enable a detailed investigation into the biological mechanisms underlying human phenotypes by integrating multiple omics, such as proteomics, metabolomics, and microbiomics [13]. Multi-omics profiling, together with clinical measures such as cytokines and blood tests, can comprehensively assess one’s health status and detect significantly correlated exposures to understand the impact of the external exposome on human biology and health [10,14,15]. In addition, longitudinal profiling can avoid biases introduced by one-time sampling and provide a molecular portrait of the effect of different exposures at an individual level.

In this first of its kind study, we utilized our previously published datasets to integrate thousands of longitudinally measured chemical and biological components along with physical factors in the personal exposome to investigate how the various stressors in the external exposome impacted internal -omes, such as the proteome, metabolome, the gut microbiome as well as cytokines and blood markers [10,14,15]. Specifically, this study 1) improved the annotation of biological and chemical exposures in the external exposome and human blood; 2) integrated the external exposome with internal multi-omics to investigate the effect of the exposome on molecular phenotypes and pathways; and 3) correlated the environmental stressors with clinical measurements to associate the health effects of the external exposome. Overall, we found thousands of external chemical and biological exposures associated with the internal microbial, proteomic, and metabolic alterations indicating a strong correlation between the external exposome and molecular health.

## Results

### Longitudinal profiling of the exposome and internal multi-omics to monitor personal environmental health

We investigated whether the external exposome is related to the internal molecular and physiology profile at a comprehensive and personal level using the schematic shown in **Figure 1a**. We first reanalyzed deep biological and chemical exposome data collected from our previously published study in which an individual had been continuously wearing a personal exposome collection device “exposometer” and correlated it with the internal molecular profiles. Over the 52-day period relevant for this study, the device captured organic chemicals using zeolite, followed by methanol elution and liquid chromatography coupled to high-resolution mass spectrometry (LC-HRMS) analysis. Biological specimens were also captured using polyethersulfone filters and nucleic acids analyzed by high throughput sequencing of DNA and RNA. (**Supplementary Data 1**). General environmental factors (e.g., temperature, humidity, total particulate matter) were recorded by the device, and the other environmental factors were also obtained from the local air quality monitoring stations (**Figure 1a**). Contrary to conventional exposome monitoring studies, which usually focused on the exposures at a single timepoint [16–18], we captured personal exposome profiles across 18 timepoints, and annotated 1,265 genera, 158 known chemical stressors among 3,299 chemical features, and 10 environmental factors which include physical stressors that may impact environmental health in this study (**Figure 1b**). These genera and known chemical stressors were annotated from improved microbiome and chemical annotation pipelines that we developed as part of this study (**Methods**).

**Figure 1.**
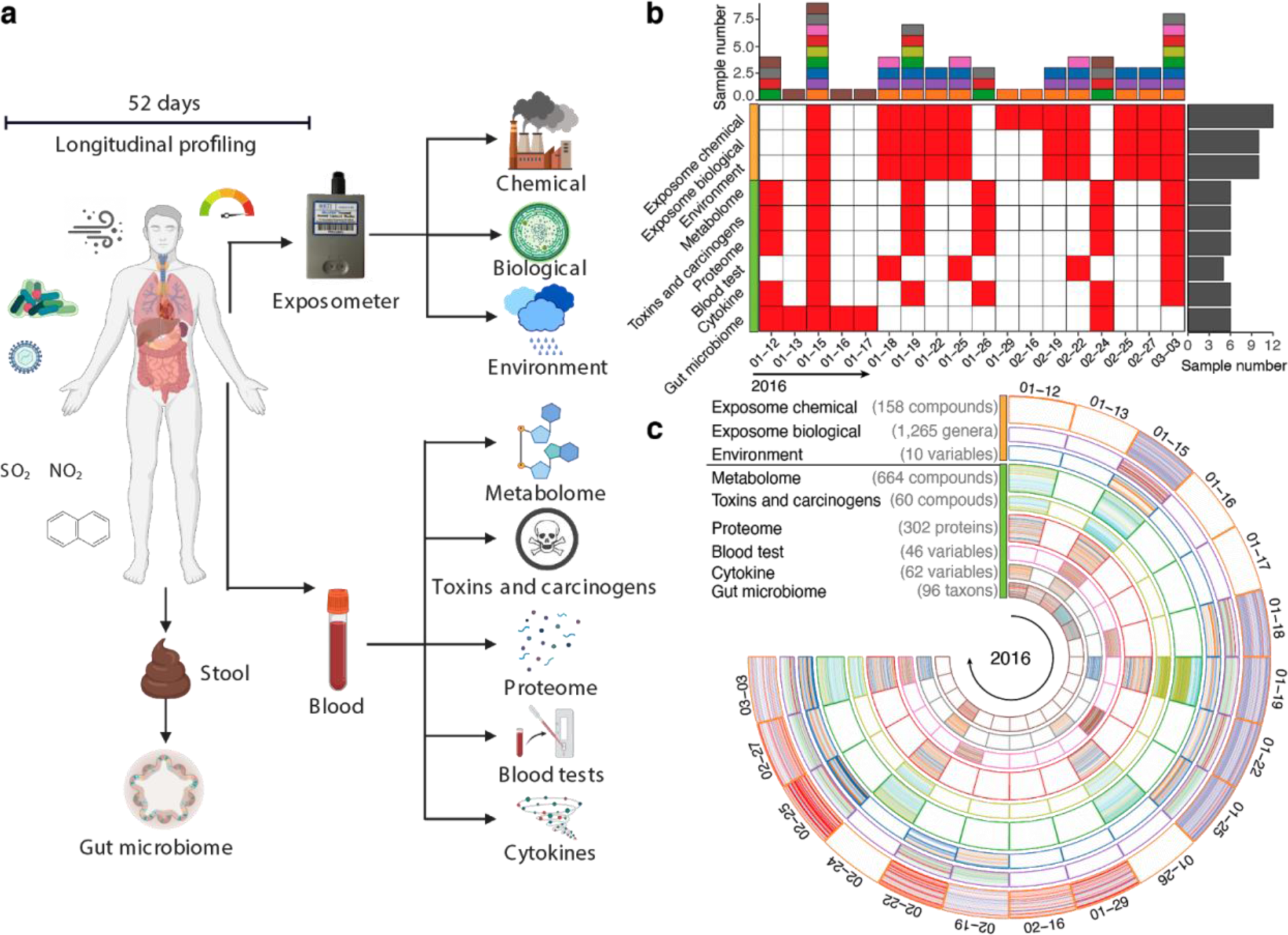
Overview of longitudinal sample collections for personal exposome and multi-omics profiling. (a) Personal exposome characterized by the exposometer includes environmental factors, biological components, and chemical stressors. Internal multi-omes include gut microbiome, metabolome, proteome, toxins and carcinogens, cytokines, and blood tests. (b) The amount and collection time of each type of multi-omics and exposome samples. (c) Sample distribution and constitution of the exposome and internal multi-omics for monitoring precision environmental health.

Over the same 52 days period, we also collected stool and blood samples from the same participant to profile the gut microbiome, proteome, metabolome, toxins and carcinogens, cytokines, and blood tests (**Figure 1c** and **Supplementary Data 1**). Through reanalysis pipelines, we were able to annotate 60 toxins and carcinogens as well as 664 metabolites, 302 proteins, and 62 gut microbiome taxa. We also measured 62 cytokines and 46 clinical blood parameters to longitudinally monitor personal health status [15]. All sample collections were performed during the first quarter of 2016 from three distinct locations in the U.S. (**Figure S1**). However, not all sample types were collected at each time point, and the inter-omics analyses were performed only when overlaps were available (**Figure 1b**). Despite our limited ability to control all confounding variables, we searched for significant intra- and inter- exposome correlations and high-degree components which have the most significant correlations in each analysis as those may play important roles in the exposome-ome interactions [|r| > 0.9; False Discovery Rate (FDR) adjusted p-value (q-value) < 0.05].

### Intra-exposome relationships in the highly dynamic and diverse personal exposome cloud

To annotate as many chemicals as possible, we searched through the 3,299 LC-HRMS raw features using a combination of five public exposome related databases as well as an in-house database that we assembled. Using this new annotation pipeline, we were able to annotate 158 known chemical stressors (**Figure 2a, Methods**). These stressors were categorized into 13 classes, with the dominant class being agrochemicals, followed by pharmaceuticals & personal care products (PPCPs), plasticizers, and International Agency for Research on Cancer (IARC) Group 2A carcinogens, and the chemicals in each class varied dynamically during the monitoring period (**Figure 2a, Figure S2** and **Supplementary Data 2**). To characterize the biological exposome domain, we circumvented the limited ability of 16S rRNA/18rRNA/ITS sequencing by applying metagenomic sequencing. We found 17 genera dominated during the study period, most of which were fungi and bacteria, but they varied dynamically (**Figure 2b**). Ten general environmental factors, measured either by personal exposometer (temperature, humidity, and total particulate matter) or local air monitoring stations (atmospheric pressure, wind speed, SO2, NO2, O3, CO, and air quality index), were also included in the study (**Figure 2c**).

**Figure 2.**
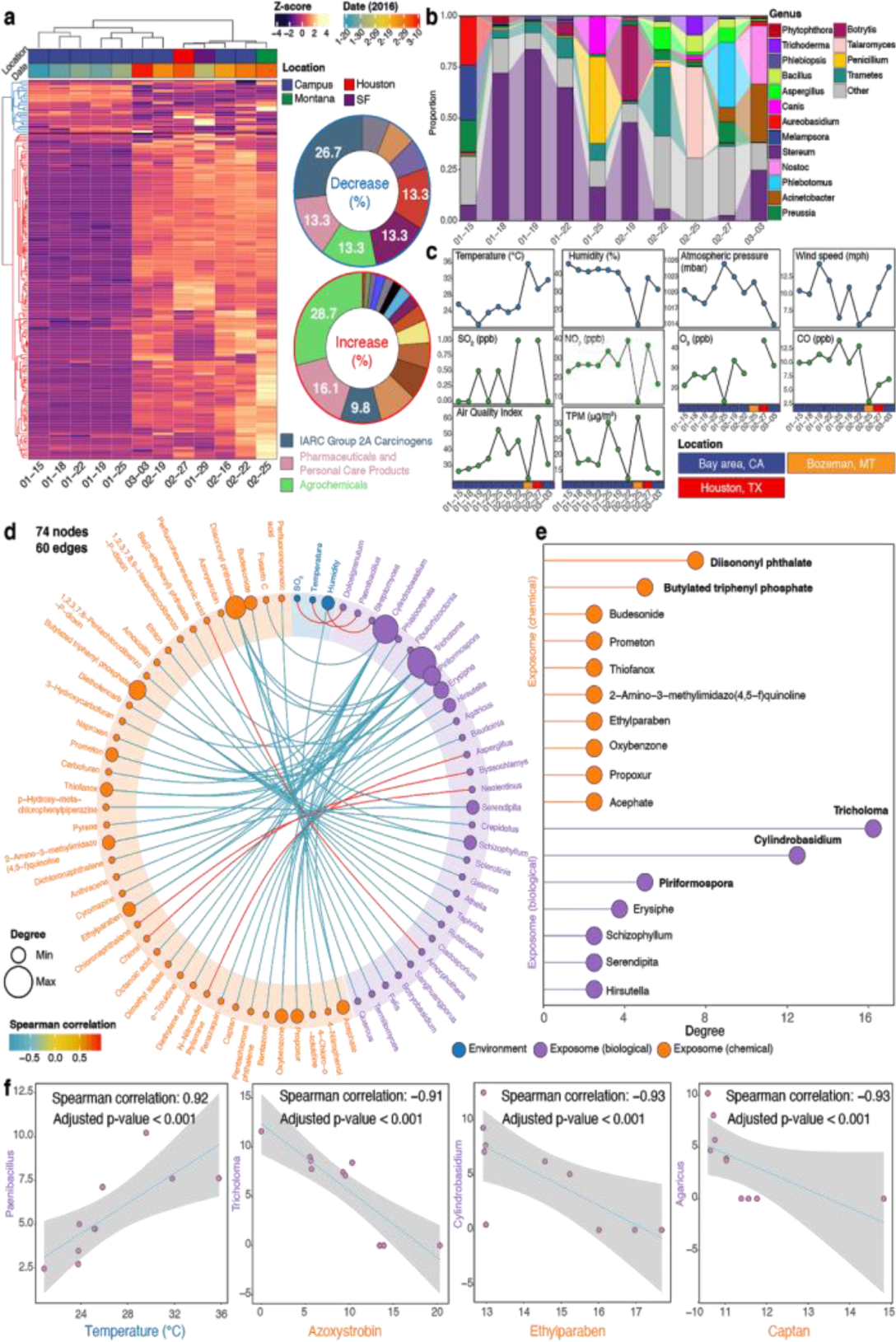
The dynamic and diverse personal exposome cloud. (a) Heat map of the annotated chemical stressors in the exposome ordered by concentrations, with the sector diagrams indicating the increased and decreased chemical groups. The abrupt concentration increase after the January 25 sample indicates the approach can monitor dramatic changes of the chemical exposome. (b) Heat map of the top abundant genera annotated in the exposome during each collection period. (c) Environmental factors collected by either the personal exposometer (temperature, humidity, total particulate matter) or local monitoring stations. TPM: total particulate matter. (d) Spearman correlation analyses within the personal exposome [|r| > 0.9; False Discovery Rate (FDR) adjusted p-value (q-value) < 0.05]. (e) Chemical and biological components that have the most significant correlations with the other substances in the exposome. (f) Representative Spearman correlation analyses between fungi and temperature/antifungal chemicals.

We performed intra-omics correlation analyses to investigate the potential relationships among all the exposome components (**Methods, Supplementary Data 3**). We found a total of 60 statistically significant correlations (|r| > 0.9 and q-value < 0.05) among 74 exposome components, including 41 chemicals, 30 genera, and 3 environmental factors (**Figure 2d** and **Figure S3**). Specifically, diisononyl phthalate (a plasticizer) and butylated triphenyl (an organophosphate flame retardant) had the most significant correlations, followed by various agrochemicals, PPCPs, and IARC group 2A carcinogen. Among the biological components, *Tricholoma* had the highest number of significant correlations, followed by *Cylindrobasidium*, *Piriformospora*, *Erysiphe*, *Schizophyllum*, *Serendipita*, and *Hirsutella*, all of which are fungi (**Figure 2e**). In terms of environmental factors, only temperature and humidity collected by the exposometer as well as SO2 concentration collected by the local monitoring stations were significantly correlated with other exposome components (**Figure 2d**). For example, *Paenibacillus* was positively correlated with the temperature, consistent with the literature that members of *Paenibacillus* are heat resistant and grow well in relative hot temperatures [19]. Azoxystrobin, ethylparaben and captan are fungicides or antifungal agents [20, 21] that negatively correlated with different fungi (**Figure 2f**).

### Inter-omics analyses between the exposome and multi-omics revealed physiological links to the exposome

To investigate how the exposome shapes an individual’s phenome longitudinally, we investigated the links between the exposome and internal multi-omics. Specifically, we found 8,986 significant correlations (|r| > 0.9 and q-value < 0.05) among 1,700 factors from all -omes, and positive correlations were more predominant than negative correlations (**Figure 3a, Supplementary Data 4**). The biological exposome and metabolome were the most extensive -omes in the network, and they also had the greatest number of significant correlations (N = 4148; **Figures 3a and 3b**). Additionally, we found that the exposome and internal multi-omics networks can be divided into several subnetworks with high modularity (0.819, **Figure S4a, b**).

**Figure 3.**
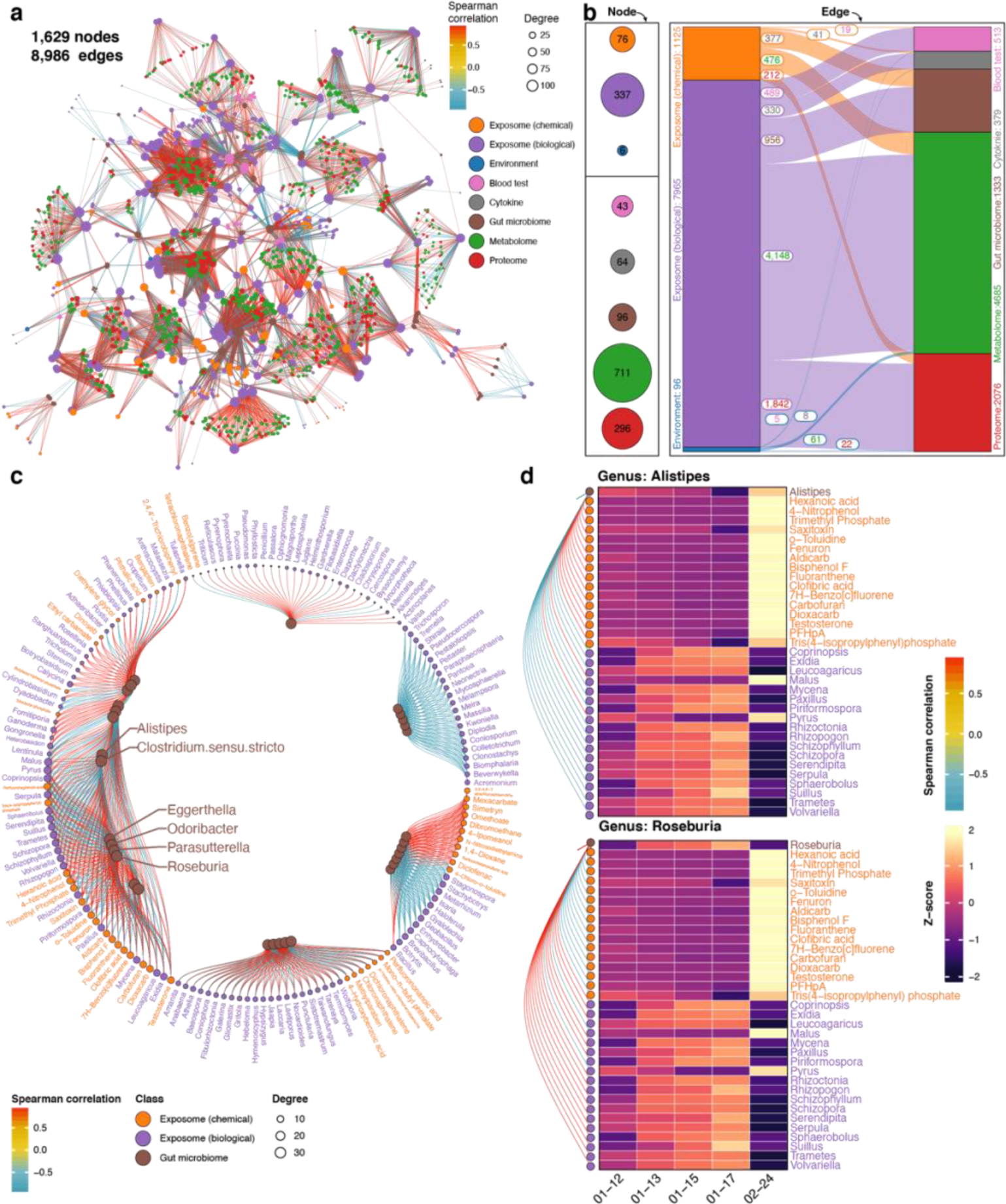
Precision environmental health network revealed by inter-omics analyses between the exposome and internal multi-omics. (a) Spearman correlation network of all longitudinally profiled exposome and internal -omes. (b) Significant Spearman correlations between the exposome and the internal multi-omics (|r| > 0.9; q-value < 0.05). (c) Spearman correlation analysis between the individual’s exposome and the gut microbiome. Only gut bacteria with degrees > 20 are shown, and the highest-degree bacteria are named. The complete network is provided in **Figure S4c**. (d) Heat map of the highest-degree personal gut bacteria and significant correlations with the exposome components (|r| > 0.9; q-value < 0.05). Additional high- degree gut bacteria are provided in **Figure S4d**.

### Personal exposome-gut microbiome interactions

We found 1,333 significant correlations (|r| > 0.9 and q-value < 0.05) between the exposome and the gut microbiome (16S rDNA data), and the number of positive and negative correlations were approximately equal (**Figure 3c**). Specifically, the six highest-degree bacteria (each correlates with 34 exposome components) may be involved in multiple physiological processes that respond to the personal exposome. For example, members from *Alistipes* were shown to play essential roles in inflammation and various diseases [22], members from *Eggerthella* were implicated as the causes of liver and anal abscesses, ulcerative colitis, and systemic bacteremia [23], members from *Odoribacter* were found to maintain short-chain fatty acid availability and systolic blood pressure [24], members from *Parasutterella* were involved in bile acid maintenance and cholesterol metabolism [25], whereas members from *Roseburia* played vital roles in producing short-chain fatty acids and anti-inflammatory pathways [26]. Out of the top six genera, all but *Roseburia* positively correlated with chemical stressors and usually negatively correlated with biological components (**Figure 3d and S4d**). As a result, members from *Alistipes*, *Eggerthella*, *Odoribacter*, and *Parasutterella* were more likely to be involved in proinflammatory processes, while members from *Roseburia* were mainly involved in anti-inflammatory processes. On the exposome side, *Botryosphaeria*, *Corynespora*, and *Enterobacter* were the highest-degree genera (each correlated with 18 gut bacteria) among all exposome components, indicating their essential roles in interacting with the participant’s gut microbiome (**Figure 3c**). Overall, these results demonstrate an association of the external exposome with the gut microbiome and its associated biological processes, particularly inflammation.

### Exposome-proteome interaction network

We found 2,054 statistically significant correlations (|r| > 0.9 and q-value < 0.05) between the individual’s exposome and internal proteome. Most of the high-degree exposome components were biological components, and positive correlations were slightly more frequent than negative correlations (**Figure 4a**). Specifically, we found 11 highest-degree substances (nine genera and two chemicals), each of which was significantly correlated with more than 22 proteins in the proteome. The high-degree biological genera were fungi and primarily positively correlated with proteins; in contrast, *Xeromyces* negatively correlated with proteins. Fenazaquin (a pesticide) and tetrabromobisphenol A diallyl ether (a brominated flame retardant) were two high-degree chemical stressors, both of which primarily negatively correlated with proteins. On the proteome side, 17 highest-degree proteins (each correlated with 21 exposome components) were discovered, and 14 of these are directly immune-related. For instance, alpha-1-HS-glycoprotein (AHSG) promotes endocytosis, complement component 3 (C3) activates the complement system, and fibrinogen alpha chain (FGA) is involved in both innate and T-cell mediated pathways [27]. Additionally, we discovered significantly correlated signaling pathways when queried against GO, KEGG, and Reactome databases (**Supplementary Data 5 and 6**). Chemical and biological exposome shared several significantly correlated pathways, such as protein activation cascade, platelet degranulation and acute inflammatory response, whereas some pathways were uniquely correlated with the chemical exposome, such as platelet activation, signaling, and aggregation pathway (**Figure 4b**). Moreover, immune-related pathways were among the most common high-degree signaling pathways correlating with chemical and biological exposome, and those pathways were about half positively and half negatively correlated with the exposome (**Figure 4c**).

**Figure 4.**
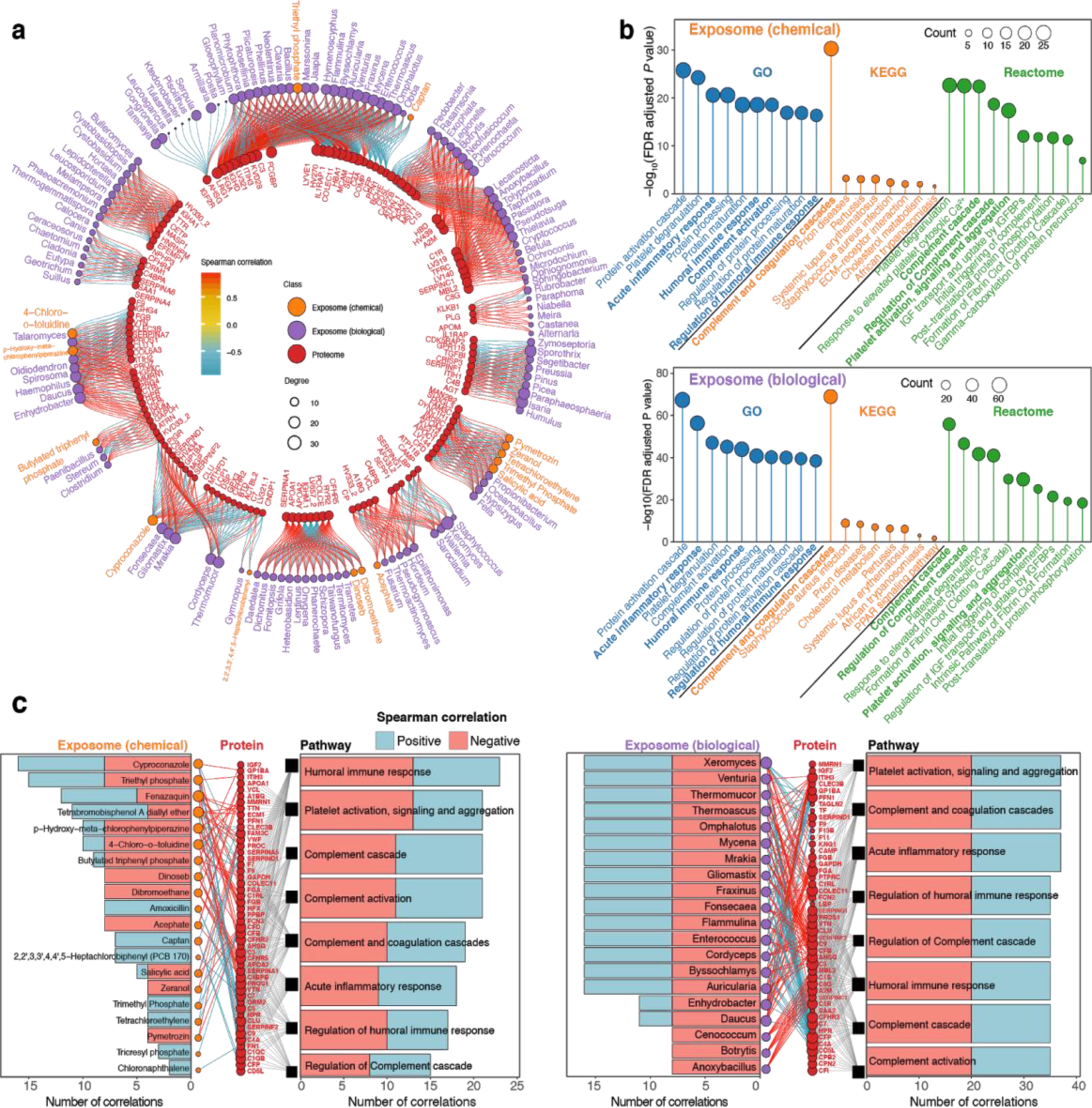
Exposome-proteome interactions: proteins and signaling pathways that were significantly correlated with the exposome. (a) Spearman correlation analysis between the exposome and proteome (|r| > 0.9; q-value < 0.05). Only the proteins with degree > 5 are shown. The complete network is provided in **Figure S5**. (b) Signaling pathways that significantly correlated with the exposome revealed by pathway analysis using KEGG, GO, and Reactome databases. Immune-related pathways are shown in bold. (c) Spearman correlation networks between chemicals, top twenty biological exposome components, immune-related proteins and signaling pathways (|r| > 0.9; q-value < 0.05), with positive correlations shown in blue and negative correlations shown in red. A detailed network for each pathway was provided in **Figure S6**.

### Exposome-metabolome interaction network

The blood metabolome is considered the most interactive -ome with the exposome since xenobiotics interact with endogenous metabolites initially after entering the human body. In fact, the blood exposome overlaps with the blood metabolome from an analytical perspective as current approaches cannot distinguish the sources of the molecules present in blood. Moreover, xenobiotic biotransformation is similar to that of metabolic pathways and can even involve the same enzymes, such as cytochromes P450 [16, 28]. Therefore, it is essential to investigate the interactions between the exposome and metabolome to better understand the initial health impact of the exposome.

In this study, we found 4,624 statistically significant correlations (|r| > 0.9 and q-value < 0.05) between the exposome and internal metabolome. Specifically, positive correlations were more frequent in the exposome-metabolome analysis than the exposome-proteome analysis (**Figure 5a and S5**). The high-degree biological components were primarily fungi and usually positively correlated with the metabolites; interesting exceptions are *Aegilops* (a grass), the bacteria *Pontibacter* and *Hymenobacter,* and *Paramecium* (a ciliated protist). Salicylic acid (a PPCP), dinoseb (an herbicide), dibromoethane (an IARC group 2A carcinogen) were the three highest-degree chemicals, all of which primarily positively correlated with endogenous metabolites. Importantly, we found 19 high-degree metabolites, each significantly correlated with 21 exposome substances. Several metabolic pathways were significantly correlated with both the chemical and biological exposome (**Methods**, **Figure S7**), such as protein digestion and absorption and aminoacyl-tRNA biosynthesis, whereas some pathways were only correlated with the biological exposome (**Figure 5b, Supplementary Data 7 and 8**). Similar to the exposome-proteome analysis, we performed correlation network analysis among the exposome, metabolites, and metabolic pathways. Trimethyl phosphate (a plasticizer and organophosphate flame retardant), 2,2’,3,3’,4,4’,5-Heptachlorobiphenyl (a polychlorinated biphenyl), and tetrachloroethylene (an IARC group 2A carcinogen) were positively correlated with all the metabolic pathways, whereas tetrabromobisphenol A diallyl ether, salicylic acid, and zeranol (a mycotoxin) were negatively correlated with all metabolic pathways (**Figure 5c**).

**Figure 5.**
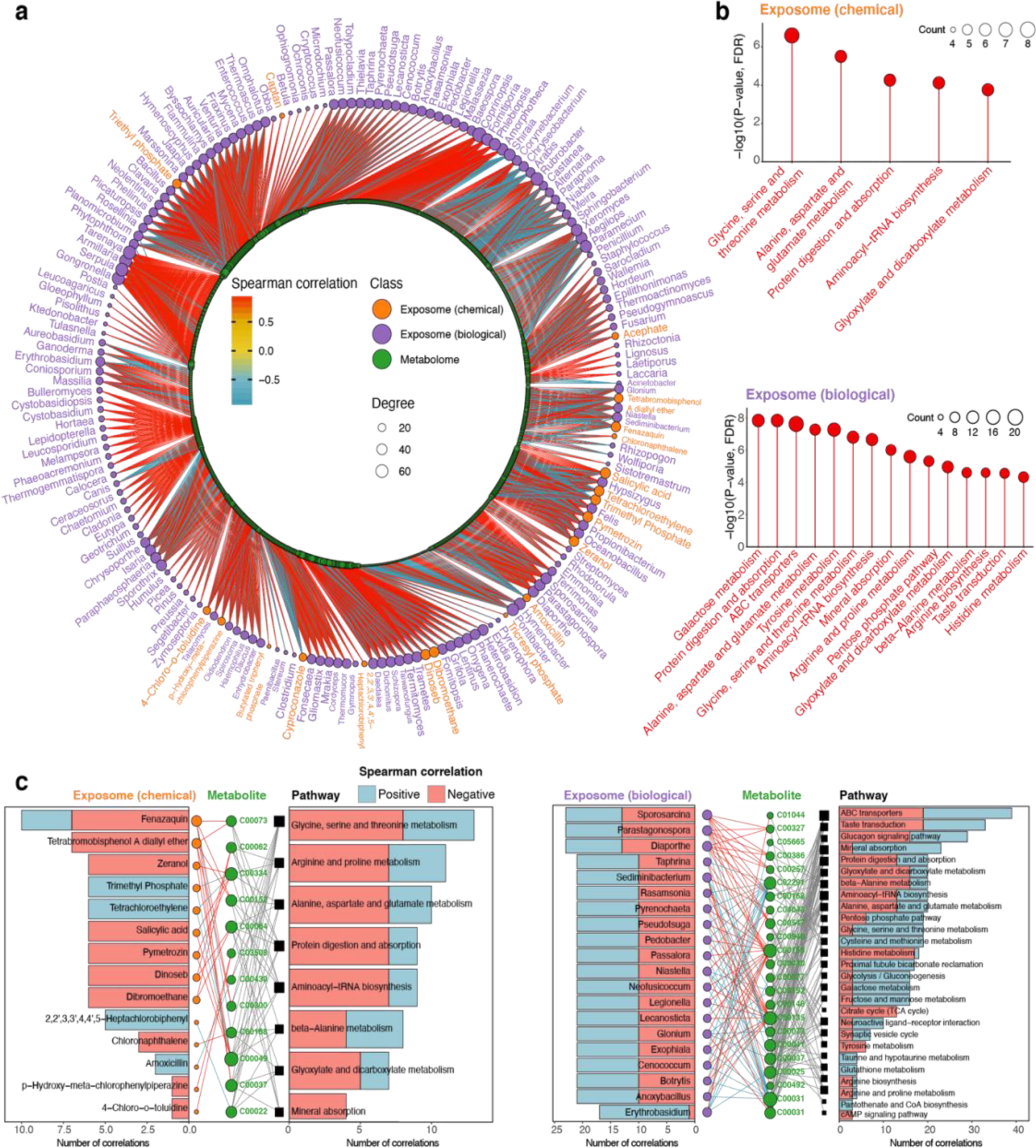
Exposome-metabolome interactions: metabolites and metabolic pathways that were significantly correlated with the exposome. (a) Spearman correlation analysis between the exposome and metabolome. (b) Significantly correlated metabolic pathways revealed by pathway analysis using KEGG database (|r| > 0.9; q-value < 0.05). (c) Significant correlations between chemicals and top twenty biological exposome components and metabolites (represented by KEGG compound entry) and metabolic pathways revealed by Spearman correlation networks (|r| > 0.9; q-value < 0.05), with positive correlations shown in blue and negative correlations shown in red. The complete network was provided in **Figure S8**.

### Monitoring precision environmental health by investigating the exposome-clinical data correlations

Standard clinical measurements such as blood and cytokine tests directly reflect the individual’s health. Thus, clinical test results are ideal indicators to investigate the health impact of the exposome. Based on our exposome-cytokine analysis, the biological exposome had the most significant correlations with cytokines, followed by chemical and environmental factors. 362 significant correlations (|r| > 0.9 and q-value < 0.05) were found between the exposome and cytokines, most of which were positive correlations. After converting correlation coefficients to variable importance in projection scores, we determined the contributions of all significantly correlated exposome components on cytokines (**Supplementary Data 10**, **Methods**). Specifically, 60% of the cytokine variation was explained by the determined factors in this study. Furthermore, the top 13 cytokines, which were almost entirely contributed by the annotated exposome components (> 90%), were all proinflammatory cytokines, such as IL-23, MCP-1, and VCAM- 1, indicating that those cytokines may play essential roles in response to the exposome. Additionally, 14 highest-degree (each correlates with > 7 exposome components) cytokines were found to be primarily positively correlated with the exposome, whereas only MCP-1 was primarily negatively correlated (**Figure 6a**). The most high-degree biological components were fungi, such as *Wallemia*, which, interestingly, are filamentous food-borne pathogens [29]. Moreover, other than *Xeromyces,* most of the exposome components were primarily positively correlated with cytokines, consistent with the exposome-proteome analysis where *Xeromyces* primarily negatively correlated the proteins. Interestingly, acephate (an insecticide) is the highest-degree chemical component, positively correlated with 10 cytokines (**Figure 6a**).

**Figure 6.**
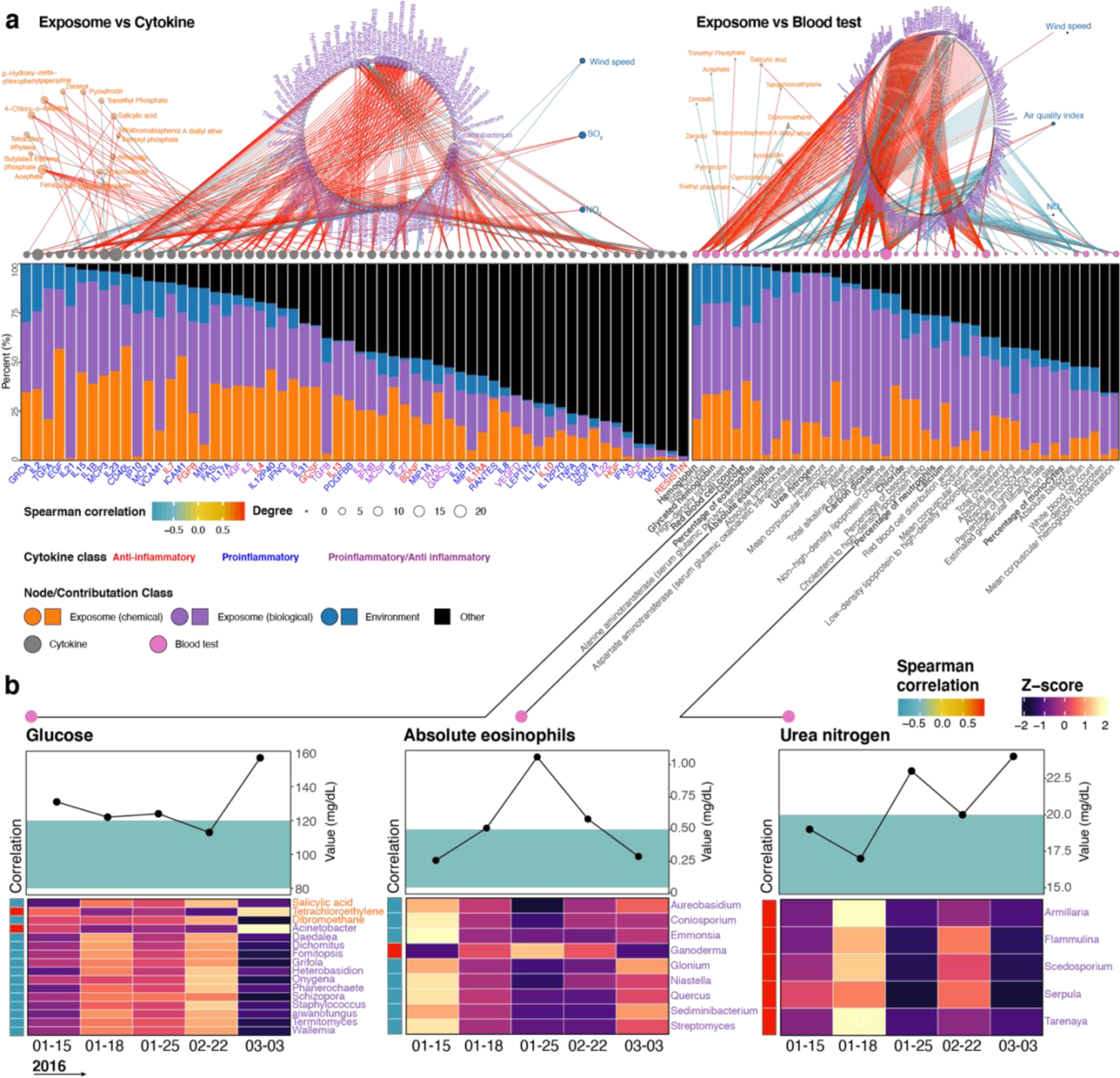
Effects of the exposome on precision environmental health. (a) Relative contributions of various exposome components on the alterations of personal cytokines and blood tests (|r| > 0.9; q-value < 0.05). (b) Representative blood test results with corresponding reference ranges (green areas) and their significantly correlated exposome components.

Similar to the exposome-cytokine analysis, biological components had the most significant correlation with blood tests, followed by chemicals and environmental factors. However, fewer chemicals were correlated with blood tests than those correlated with cytokines (**Figure 6a**). 513 significant correlations were found between the exposome and blood tests, and the majority were positive correlations. Using similar contribution determination algorithms, 77% of the blood tests variation was explained by the determined factors in this study. Similarly, the top 13 blood tests which were almost entirely contributed by the determined exposome components (contributions of the exposome > 95%), were primarily related to the immune system, liver, and kidney functions. Additionally, eight highest-degree (each correlated with > 25 exposome components) blood tests were primarily positively correlated with the exposome while only platelet was primarily negatively correlated. Interestingly, the highest-degree blood test, creatinine, which is a biomarker for kidney function, correlated with 62 exposome components (**Figure 6b**). Unlike cytokine profiles, where we cannot draw a clear line of the individual’s health status, blood tests have clinically established reference ranges facilitating the clinical impacts. We therefore performed correlation analyses to understand the effects of the exposome on personal health using blood test results with out-of-range values. Interestingly, we found the abnormal blood glucose level values were significantly correlated with 3 chemical stressors and 13 microbes. For instance, salicylic acid concentration was negatively correlated with glucose level; salicylic acid has been previously shown to decrease glucose concentration and used as a treatment for type 2 diabetic patients [30], which is consistent with our findings. Similarly, abnormal values of absolute eosinophils and urea nitrogen correlated with specific biological exposome components, including some known pathogens, such as *Aureobasidium*, *Niastella*, and *Scedosporium* (**Figure 6b**). Previous studies were consistent with our results as eosinophilic phagocytosis consumes eosinophils during allergy and inflammation [31], and various pathogenic microbes can utilize urea as a nitrogen source [32].

## Discussion

It has long been acknowledged that environmental factors affect personal health, but conventional environmental health studies face limitations. For example, a) population or cohort studies overlook the significant differences between individuals; b) single timepoint sampling fails to reflect the continuous effects of stressors; c) and focusing on a single or a class of stressors does not capture the holistic health impact of the exposome. To overcome these challenges, we generated a comprehensive precision environmental health profile by longitudinally monitoring both the personal exposome and internal multi-omic profiles (**Figure 1**). In addition, we also measured standard clinical indices to investigate the health effects of the exposome. Using Spearman correlation analysis, we discovered many significant correlated physiological parameters and exposome components, indicating their interactions in the participant’s responses to the personal exposome. Additionally, our study provided vast testable hypotheses to further investigate the underlying mechanisms using analytical and experimental approaches.

We were able to capture more than chemical 3,000 features, but only annotated 158 known chemical stressors by a broad annotation method that utilizes various databases, including those containing emerging contaminants [33–36]. This indicates that existing exposome databases still lack the power to annotate the majority of the chemical exposome. Interestingly, we found that the concentrations of most chemicals increased after January 25, 2016, when the individual transitioned from a period of residing at home to a period of high travel, indicating that the chemical exposome greatly increased with travel to other locations (**Figure S2f**). Agrochemicals had the highest concentrations among all annotated chemicals, indicating their ubiquitous presence in the environment. An alternative view is that agrochemicals are the most frequently studied chemical stressors, making them most easily identifiable. It is also worth noting that high concentrations do not necessarily imply high health risks since each chemical has its own safe dose, and the combined effects among them are still unclear [13].

The biological exposome revealed a number of interesting observations as well. The fungal genus *Stereum* was dominant at most time points, reflecting its high abundance in the personal exposome (**Figure 2a**). We found several interesting correlations of the biological exposome with chemical and environmental factors, such as associations of antifungals with a decrease in fungal exposures. Many of these are intended food or soil antifungal products (azoxystrobin and captan), whereas others are common preservatives (ethylparaben and anthracene) that have antifungal properties [10, 11]. Overall, nearly 100 significant correlations were found by intra-exposome analysis, representing the complex interactions within the exposome domains.

Importantly, the negative correlations of fungi with various pesticides and herbicides indicate these agrochemicals may inhibit the fungi growth as well (e.g., *Tricholoma* versus propoxur and *Erysiphe* versus bentazone). Moreover, we find several interesting tertiary relationships, such as the mycotoxin fusarin C (produced by *Fusarium*) negatively correlated with *Cylindrobasidium,* suggesting a possible competition among the different fungi (**Supplementary Data 3**).

A recent study identified radioprotective gut microbes and internal metabolites in mice using a multi-omics analysis [37], demonstrating the potential of this approach to investigate essential components in the internal -omes that respond to the external environment. To this end, we performed inter-omics analyses between the exposome and gut microbiome, proteome, and metabolome, respectively. By discovering high-degree components in each analysis, we identified the critical components in the exposome-internal omes interactions. For instance, we found six highest-degree gut bacteria that may be important in the responses to the personal exposome. The high-degree biological components in both exposome-proteome and exposome-metabolome analyses were mainly fungi, yet few had known human health effects. However, we identified major high-degree annotated chemicals that are known human stressors; for instance, the herbicide dinoseb exposure causes various developmental toxicities and loss of thyroid and body weight [38], and brominated flame retardants like tetrabromobisphenol A diallyl ether are known neurotoxicants [39]. In addition to the endogenous metabolites, we were able to annotate 60 toxins and carcinogens in the individual’s blood based on the exposome related databases (**Figure 1c**). Unlike the annotated xenobiotics in the exposome samples, most of the chemical stressors annotated in the blood were food and animal toxins. Furthermore, only 11 chemical stressors were annotated in both the external and blood exposome. This is likely to be partly due to the limited power of the current databases, since most of the databases only contain the information of parent chemicals but not their biotransformation products. Additionally, persistent hydrophobic substances tend to accumulate in adipose tissues, but not in blood (which we profiled), while non-persistent hydrophilic chemicals are efficiently excreted out of the human body [13, 16], limiting their detections. Finally, the bioavailability of chemicals in different external matrices also limits the exposure, dose-response, and concentration of bioavailable fraction [35,40,41].

Importantly, our results indicate that the immune system, kidneys, and liver may play essential roles in response to the exposome, which are all known to regulate and respond to foreign substances [17,42,43]. In the exposome-proteome analysis, we found 14 out of 17 high-degree proteins were involved in immune responses (e.g., complement component 3, interleukin-1 receptor accessory protein, and immunoglobulin heavy chain proteins) [43], and immune-related pathways (e.g., acute inflammatory response, humoral immune response, and complement and coagulation cascades) and were among the highest-degree signaling pathways. In the exposome-metabolome analysis, we identified 19 highest-degree metabolites related to protein metabolism, inflammation, kidney and liver functions (e.g., L-arginine, nutriacholic acid, epsilon- (gamma-Glutamyl)-lysine, and uracil), indicating that these metabolic pathways were involved in responses to the exposome. Moreover, certain high-degree metabolic pathways are both protein and immune-related pathways, such as alanine aspartate and glutamate metabolism, protein digestion and absorption, and beta-alanine metabolism. Specifically, particular protein metabolism pathways (e.g., amino acids synthesis and protein breakdown) were highly sensitive to oxidative stresses caused by the exposome components [44], inflammation is often the first immunological response to foreign substances, and kidneys and liver are the main detoxification organs [44], with liver and bile acids serving essential roles in responding to the foreign substances [45].

To further investigate the health effects of the exposome, we performed Spearman correlation analysis with cytokines and blood test results. Proinflammatory cytokines were the most significantly correlated with the external exposome components (e.g., IL-23, MCP-1, and IL-2), and they have been previously shown to be elevated after exposure to external stressors [46]. Our blood test results provided further evidence as fluctuations of creatinine and urea nitrogen, biomarkers of kidney and liver functions, respectively, were correlated with specific exposome components. As a result, exposome-proteome analysis cross-validated with exposome-cytokine analysis, indicating that the proinflammatory processes play essential roles in responding to the exposome, and the exposome-metabolome analysis cross-validated with exposome-blood tests analysis, showing that liver and kidneys play significant roles in responding to the exposome. Furthermore, the exposome-microbiome analysis showed that the highest-degree gut bacteria are related to the proinflammatory processes and liver metabolism. Therefore, these physiological processes and organs may be ideal candidates for testing the combined effects of multiple stressors in future studies.

On the exposome side, high-degree exposome components that overlapped in more than one inter-omics analysis are significant health concerns. Specifically, *Isaria*, *Sporothrix*, and *Tarenaya* were among the highest-degree microbes correlated with all internal -omes. Members of these genera were found to be involved in complex physiological mechanisms and may exhibit adverse health effects. For example, species of *Isaria* were found to induce cell death [47], species of *Sporothrix* triggered skin and lung inflammatory reactions [48], and the pollen of *Tarenaya* members are allergens and generate immunoglobulin-E mediated allergic reactions [49]. Also, certain types of agrochemicals, PPCPs, and flame retardants that we detected are high-degree chemical stressors known to cause epigenetic alterations, endocrine disruption, impaired nervous system function, oxidative stress, and inflammation [44]. Therefore, those substances would be ideal candidates for investigating the underlying mechanisms of their combined health effects in future studies. Such information may be especially important for understanding environmental triggers for inflammatory diseases such as inflammatory bowel diseases, autoimmune diseases, and skin inflammation.

In conclusion, we found both time and location impacted the personal exposome, especially the biological components and environmental factors (**Figure S2f**), and the biological and chemical exposome was highly dynamic. These results emphasize the significance of individual precision environmental health over traditional environmental epidemiology studies. Undoubtedly, due to the limitation of the single participant and annotation power in this study, future research should monitor the precision environmental health of more individuals and increase the annotation confidence of the chemical and biological components in the exposome. In addition, we focused on the airborne exposome and did not measure other exposures, such as dermal and ingestion exposures, inorganic chemical components [50–53], psychosocial stressors, as well as personal lifestyle, which may affect the clinical measurements as well (**Figure 6a**). Nonetheless, this study demonstrates the power of using a holistic approach of monitoring the exposome on personal environmental health using inter-omics analyses, and serves as a useful model to scale to the other individuals and locations. Our study also identified high-degree components as essential components among the exposome-internal omes interactions and provided abundant testable hypotheses to further investigate their underlying mechanisms of impacting individual health.

## Methods

### Personal exposome and internal multi-omics samples collection Exposome samples collection

The participant in the study is enrolled under Stanford University’s IRB protocols IRB-23602 and IRB- 34907. Modified RTI MicroPEM V3.2 personal exposure monitor (RTI International, Research Triangle Park, North Carolina, USA), termed exposometer, was used to collect chemical and biological components exposed by individuals from January 2016 to March 2016. Also, temperature, humidity, and total particulate matter were simultaneously collected by the exposometer in a real-time manner. The original sequential oiled frit impactor was removed to maximize the collection of biological components. A 0.8 mm pore-size polyethersulfone with a diameter of 25 mm filter (Sterlitech, Kent, Washington, USA) was placed in filter cassettes to collect particulates for DNA and RNA extraction. An in-house designed, 3D printed cartridge was placed at the end of the airflow, which contained 200 mg of zeolite adsorbent beads (Sigma 2-0304) to collect chemicals. Before deployment to the participant, the MicroPEM was calibrated to a flow rate of 0.5 L/min (± 5%) using a mass flow meter (TSI 4140, Shoreview, Minnesota, USA). During the study, the participant was instructed to place the exposometer on his arm or within a radius of 2 meters. Samples were collected after 1 to 3 days of use and stored at -80 °C until analysis. To minimize the potential contamination, filters and related components were handled in sterile biological safety cabinets and cleaned by ethanol before use. Clean polyethersulfone filter and zeolite adsorbent beads were included before extraction as background controls. MicroPEM log files were downloaded using Docking Station software (RTI International, Research Triangle Park, North Carolina, USA). The participant used the MOVES App to track geographic locations through GPS coordinates and daily activities [10]. General environmental data were collected from the exposometer or National Oceanic and Atmospheric Administration’s National Climatic Data Center or National Centers for Environmental Information.

### Analysis of chemical exposome by LC-HRMS

LC-HRMS was performed on a platform composed of a Waters UPLC coupled with Exactive Orbitrap mass spectrometer (Thermo, Waltham, Massachusetts, USA) using a mixed-mode OPD2 HP-4B column (4.6 mm x 50 mm) with a guard column (4.6 mm x 10 mm; Shodex, Showa Denko, Tokyo, Japan). The column temperature was maintained at 45 °C and the sample chamber at 4 °C. The binary mobile phase solvents were: A, 10 mM ammonium acetate in 50:50 (vol/vol) acetonitrile: water; B, 10 mM ammonium acetate in 90:10 (vol/vol) acetonitrile: water. Both solvents were modified with 10 mM acetic acid (pH 4.75) for positive mode acquisition or 10 mM ammonium acetate (pH 9.25) for negative mode. The flow rate was set as follows: flow rate, 0.1 ml/min; gradient, 0–15 min, 99% A, 15–18 min, 99% to 1% A; 18– 24 min, 1% A; 24–25 min, 1% to 99% A; 25–30 min, 99% A. The MS acquisition was set as full scan mode with an ESI probe or APCI probe. The capillary temperature was 275 °C, the sheath gas was 40 units, the positive mode spray voltage was 3.5 kV, and 3.1 kV for the negative mode. The capillary voltage was 30 V, the tube lens voltage was 120 V, and the skimmer voltage was 20 V. The mass spectrum scan used 100,000 mass resolution, high dynamic range for AGC target, maximum injection time of 500 ms, and a scan range of 70-1,000 m/z. The details of quality assurance and quality control were described in the previous study [54].

### Post-acquisition analysis of the chemical exposome

Analysis of chemical exposome was performed as previously described [10, 54]. In brief, feature detection was performed with XCMS. For a conservative assessment of the number of unique chemical features, a customized Python script was used to remove potential isoforms, isotopes, and adducts from the 3,299 features enriched at least 10-fold as compared with the blank control. The annotation was based on various exposome related databases that are publicly available as well as in-house databases [55–57]. The annotation confidence levels of all the chemicals in the exposome were at least level 5, with at least one chemical at level 1 or 2 in each chemical class [54, 58].

### Sequencing and analysis of biological exposome

DNA and RNA sequencing and analysis were performed as previously described [10, 54]. In brief, DNA and RNA were extracted from filters and linearly amplified for sequencing. Libraries were sequenced by Illumina HighSeq 4000 platform (2 x 151bp) that yields an average of ∼50 M unique reads per sample. Sequenced reads were deduplicated, and adapters were trimmed using Trim Galore! (version 0.4.4). Human related reads were identified using BWA mapped to the hg19 human genome and removed. Following dehumanization, non-human reads were used for de-novo assembly using Megahit (1.1.1), and contigs were queried against our in-house database with BLASTN (2.3.9+) wrapper. The extensive in-house reference genome database included more than 40,000 species covering all domains of life [10]. Taxonomy classification and abundance were determined using a customized Lowest Common Ancestor (LCA) algorithm.

### Blood sample collection

At the designated time point, blood was drawn from the overnight fasted participant in the Clinical and Translational Research Unit at Stanford University. Aliquots of blood were condensed at room temperature to coagulate, and clots were subsequently pelleted. The serum supernatant was then immediately frozen at -80 °C. The blood in the EDTA tubes was immediately layered onto the Ficoll medium and spun with gradient centrifugation. Then the top layers were removed, and plasma was aliquoted and immediately frozen at –80 °C. Subsequently, blood mononuclear cells (PBMCs) were collected and counted using a cell counter. Aliquots of PBMCs were further pelleted and frozen with DMSO/FBS. For the later multi-omics analyses, PBMCs were thawed on ice and then lysed to protein fraction using Allprep Spin Columns (Qiagen) according to the manufacturer’s instructions with the QIAshredder lysis option. Upon receipt of samples, blood samples were then stored at –80 °C for clinical tests. The details of the blood and cytokines tests can be found in **Supplementary Data 1** [15].

### Collection and analysis of the gut microbiome

Stool samples were collected according to the Human Microbiome Project-Core Microbiome Sampling Protocol A (https://www.hmpdacc.org/). Following the Human Microbiome Project-Core Microbiome Sampling Protocol A (HMP Protocol #07-001, v12.0), DNA extraction was performed. We used the MOBIO PowerSoil DNA extraction kit and proteinase K to isolate DNA in a clean fume hood. Samples were then treated with lysozyme and staphylococcal hemolysin. For 16S (bacterial) rRNA gene amplification, primers 27F and 534R (27F: 5′-AGAGTTTGATCCTGGCTCAG-3′ and 534R: 5′- ATTACCGCGGCTGCTGG-3′) were used to amplify the 16S hyper-variable regions V1–V3. Unique barcode amplicons were used, and samples were sequenced on the Illumina MiSeq platform (V3; 2 × 300 bp). Illumina software handled the initial processing of all raw sequencing data. Reads were further processed by removing low-quality (average quality <35) and ambiguous base (Ns) sequences. UChime was used to remove chimeric amplicons, cluster the amplicon sequences, and select the operational taxonomic unit by Usearch based on the GreenGenes database (version in May 2013). The final biological classification assignment was performed using the RDP-classifier in QIIME with custom scripts [14].

### Untargeted proteomics by LC-HRMS

Preparation and analysis of plasma samples were performed as previously described [14]. In short, tryptic peptides from plasma samples were separated on the NanoLC 425 system (SCIEX). 0.5 × 10 mm ChromXP (SCIEX) was used for trap-elution settings, and the flow rate was set to 5 µl/min. The LC gradient was a 43-minute gradient with mobile phase A: 0.1% formic acid in 100% water and mobile phase B: 0.1% formic acid in 100% acetonitrile. During the gradient, mobile phase B was 4–32%. Then 8 µg of undepleted plasma were loaded on LC. SWATH acquisition was performed on a TripleTOF 6600 system equipped with a DuoSpray source and a 25 µm ID electrode (SCIEX). The variable Q1 window SWATH acquisition mode (100 windows) was constructed in the high-sensitivity MS^2^ mode. PyProphet [59] was used to score the peak groups in each run statistically, and TRIC [60] was used to align all runs. Finally, a matrix with a peptide level of 1% FDR and a protein level of 10% FDR was generated for subsequent analysis. The protein abundances were the sum of the first three most abundant peptides. Perseus (v 1.4.2.40) was applied to subtract the main components showing the main batch deviation to reduce batch effect [14].

### Cytokine profiling

The levels of circulating cytokines in the blood were measured by the 62-plex Luminex antibody-conjugated magnetic bead capture assay (Affymetrix), which has been extensively characterized and benchmarked by the Stanford Human Immunological Monitoring Center. The human 62-plex (eBiosciences/Affymetrix) was utilized with the modifications described below. Briefly, beads were added to a 96-well plate and washed using Biotek ELx405. Samples were added to the plate containing mixed antibody-linked beads, incubated at room temperature for 1 hour, and then overnight at 4 °C with shaking (500-600 r.p.m, orbital shaker). After overnight incubation, the plate was washed, and then the biotinylated antibody was added. The plate was incubated at room temperature with shaking for 75 minutes. The plate was washed, and streptavidin-PE was added for detection. After incubating for 30 minutes at room temperature, the plate was washed once, and then the reading buffer was added to the wells. The plate was read by a Luminex 200 instrument, and the lower limit of each cytokine per sample was set to 50 beads. Radix Biosolutions custom assay control beads were added to all wells. The batch effect was corrected using replicates and controls shared between batches [14].

### Untargeted metabolomics by LC–HRMS

All blood samples were prepared and analyzed for metabolomics as previously described [61]. In short, plasma samples were extracted with acetone: acetonitrile: methanol (1:1:1 vol/vol/vol), evaporated to dryness under nitrogen and reconstituted in methanol: water (1:1 vol/vol) for LC-HRMS analysis. HILIC and RPLC separations were used to analyze the extractants four times in positive and negative modes, respectively. HILIC metabolomics data was obtained on a Thermo Q Exactive plus, and RPLC metabolomics data was obtained on a Thermo Q Exactive. Both instruments were equipped with HESI-II probes and operated in the full MS scan mode. We only used the combined quality control samples from the study to obtain MS^2^ data. We used a ZIC-HILIC column (2.1 × 100 mm, 3.5 µm, 200Å; Merck Millipore) and mobile phases composed of 10 mM ammonium acetate in acetonitrile: water (50:50 vol/vol) and 10 mM ammonium acetate in acetonitrile/water (95:5 vol/vol) (B), and a Zorbax SB-Aq column (2.1 × 50 mm, 1.7 µm, 100Å; Agilent Technologies) and mobile phases composed of 0.06% acetic acid in water (A) and 0.06% acetic acid in methanol (B) to perform HILIC and RPLC analyses, respectively. All raw metabolomics data were processed using Progenesis QI (Nonlinear Dynamics, Waters). We also removed features that did not show sufficient linearity when diluted. Only features presented in more than ⅓ samples were retained for further analysis, and the KNN method was used to estimate missing values. To normalize the data, locally assessed scatter plot smoothness analysis was applied [62]. Metabolic signatures were identified by matching retention time and fragmentation spectra to corresponding standards or comparing fragmentation patterns to public repositories, as previously reported [14]. Toxin and carcinogens were annotated out of the metabolome features if the feature cannot be annotated as a human metabolite. The annotations of toxins and carcinogens were based on various blood exposome related databases that are publicly available as well as in-house databases [55–57]. The confidence levels of blood chemical annotation were at least level 5, with the majority at least level 3 [58].

### General statistical analysis and data visualization

All statistical analysis and data visualization were performed using R (v3.6.0, https://www.r-project.org/) and RStudio (v 1.2.5019). Most of the R packages and their dependencies used in this study were deployed in CRAN (https://cran.r-project.org/) or Bioconductor (https://bioconductor.org/), and some of them are deployed on Github. Session information for this study is provided in **Supplementary Note 1**. All scripts to reproduce analysis and data visualization for this study is available on Github (https://github.com/jaspershen/precision_exposome/tree/main/R/20200511). All data from the exposome and internal -omes data were log2 transformed before analysis. According to the participant’s food log, fiber intake was adjusted for all internal -omes data to reduce fiber intake biases (**Supplementary Note 2**).

### Exposome and internal multi-omics correlation networks

Spearman correlation was used to build the correlations in the intra/inter-omics analyses. In general, for each two -omes pair, the correlation matrix was calculated as below: for each variable in one -omes, Spearman correlations and FDR adjusted p-values were generated with all features in the other -omes. Only correlations between each pair variable with absolute correlation > 0.9 and FDR adjusted *p*-value < 0.05 were kept to construct the final correlation networks.

### Community analysis

Community analysis was performed based on edge betweenness embedded in R package **igraph** (https://igraph.org/). Briefly, this is an iterative process, the edges with the highest edge betweenness score were removed in each iteration, and the process was repeated until only individual nodes remain. At each iteration, modularity was calculated, and communities were analyzed at the iteration that maximized this quantity. A visualization of iteration community versus modularity is shown in **Figure S6 a, b**. To ensure the robustness and reliability of our findings, only communities (or clusters) with at least 3 nodes were kept for subsequent analysis. All the networks were visualized using R package **igraph**, **ggraph** and **tidygraph**.

### GO, KEGG and Reactome pathway enrichment for proteome

The R package **ClusterProfiler** (v 3.18.0, https://bioconductor.org/packages/release/bioc/html/clusterProfiler.html) was used for GO, KEGG and Reactome pathway enrichment for proteomics. In general, UNIPROT and ENTREZID were obtained for proteins that connect with the exposome in correlation networks. Then the GO, KEGG and Reactome pathway databases were used for pathway enrichment (hypergeometric distribution test, *p*-values are adjusted by the FDR method, and the cutoff was set as 0.05). Only pathways with hitting protein number > 3 were retained for subsequent analysis.

### Metabolic feature based dysregulated module detection

Applying the same concept from mummichog [63] and PIUMet [64], metabolic networks from KEGG and community analysis were utilized to detect dysregulated modules based on metabolic features connecting the exposome, respectively [63]. In general, the metabolic network (MN) was downloaded from KEGG, which contains 1,377 nodes (metabolites) and 1561 edges (reactions). The brief workflow is described below:

1. All the metabolic features connecting the exposome (Lsig) were matched with the KEGG metabolite database based on different adducts (**Supplementary Table 1**). Then all matched metabolites (significant metabolites) are mapped in the metabolic network to get the subnetwork (SN). Non-significant metabolites (hidden metabolites) that can connect significant metabolites within 3 reactions were also included in the subnetwork. Then the modules (M) were detected in the subnetwork via random walks [65]. Only modules with at least 3 nodes were kept. These modules were named significant modules (Msig) from real biological-related metabolic features.
2. For each module, the activity score (S) was calculated to measure both the modularity and enrichment of input metabolites (I). Activity score (S) of the module (M) was calculated as follows: For a module M: 

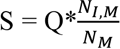
3. Where S is the activity score, NM is the metabolite number in module M, and NI,M is the input metabolite number in module M. Q is the adjusted Newman-Girvan modularity calculated as below: 

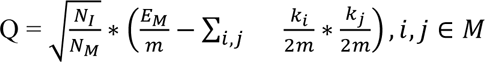 where ki is the degree of metabolite i in module M, m is the total number of edges in the metabolic network MN, EM is the total number of edges in module M, and NI is the number of input metabolites. The original Newman-Girvan modularity has a bias towards larger modules, and 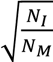 was used to reduce this bias.
4. Then the NULL metabolic features (Lnull, the same number with Lsig) were selected from all metabolic features (exclude Lsig) and then steps 1 and 2 were repeated 100 times to generate a list of NULL modules (Mnull) and their activity score (Snull).
5. Using maximum likelihood estimation, Snull was modeled as a Gamma distribution, and a cumulative distribution function (CDF) was calculated. The p-value for each significant module was then calculated, and only modules with the p-value < 0.05 remained.

The annotation results from this method were also compared with the annotation results from the “**Untargeted metabolomics by LC-HRMS**” section, provided in **Supplementary Data 9**. These results showed that annotations from this method have high specificity.

### KEGG pathway enrichment analysis for metabolomics data

The KEGG pathway database was downloaded from KEGG (https://www.genome.jp/kegg/) using the R package **KEGGREST**. Pathway enrichment analysis was used in the hypergeometric distribution test, and *p*-values were adjusted by the FDR method, and only pathways with FDR adjusted p-value < 0.05 were kept.

### Exposome contributions to cytokine and blood test

To calculate the contributions of the exposome on each cytokine and blood test, principal components (PCs) were first extracted for each exposome component, and only PCs with cumulative explained variation > 80% were kept. Then the linear regression model was constructed using each cytokine/blood test as y and corresponding exposome component’s PCs as x. R^2^ was extracted and used to represent the contributions of the exposome to each cytokine/blood test. To calculate the contribution of the exposome components, partial least squares (PLS) and variable important projection (VIP) were calculated. Finally, 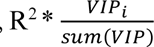 ∈ chemical, biological, and environment) was used to represent the contributions of the exposome components on cytokine/blood tests.

### Data availability

The biological exposome data generated in this and previous studies were deposited to the National Center of Biotechnology Information under bioproject ID PRJNA421162. Some raw data utilized in this study are presented on the NIH Human Microbiome 2 project site (https://portal.hmpdacc.org). Some other raw and processed data are shown on the Stanford iPOP site (http://med.stanford.edu/ipop.html). The processed data used for reproductive analysis can also be found on Github (https://github.com/jaspershen/precision_exposome) and were provided in **Supplementary Data 1**.

### Code availability

All codes used in this study can be found on Github (https://github.com/jaspershen/precision_exposome). Certain in-house tools for this study can also be found on Github (https://github.com/jaspershen/metID).

## Acknowledgements

We thank Xin Wang and Xiyan Li for their contributions to generate the raw data of chemical exposome. This work was supported by Leona M. and Harry B. Helmsley Charitable Trust (grant G-2004-03820). S.M.S.-F.R. was supported by an NIH Career Development Award no. K08ES028825. This work used supercomputing resources provided by the Stanford Genetics Bioinformatics Service Center, supported by NIH S10 Instrumentation Grant S10OD023452.

## Author contributions

P.G. and M.S. conceived the study. P.G., X.S., and C.J. analyzed the chemical exposome and environmental factors data. P.G., X.S., X.Z., and C.J. analyzed the biological exposome data. P.G., X.S., and X.Z. analyzed multi-omics data. M.S. supervised the study and provided funding support. P.G. drafted and revised the manuscript. X.S., X.Z., C.J., S.Z., X.Z., S.M.S.-F.R., and M.S. revised the manuscript.

## Supplementary information

**Figure S1.**
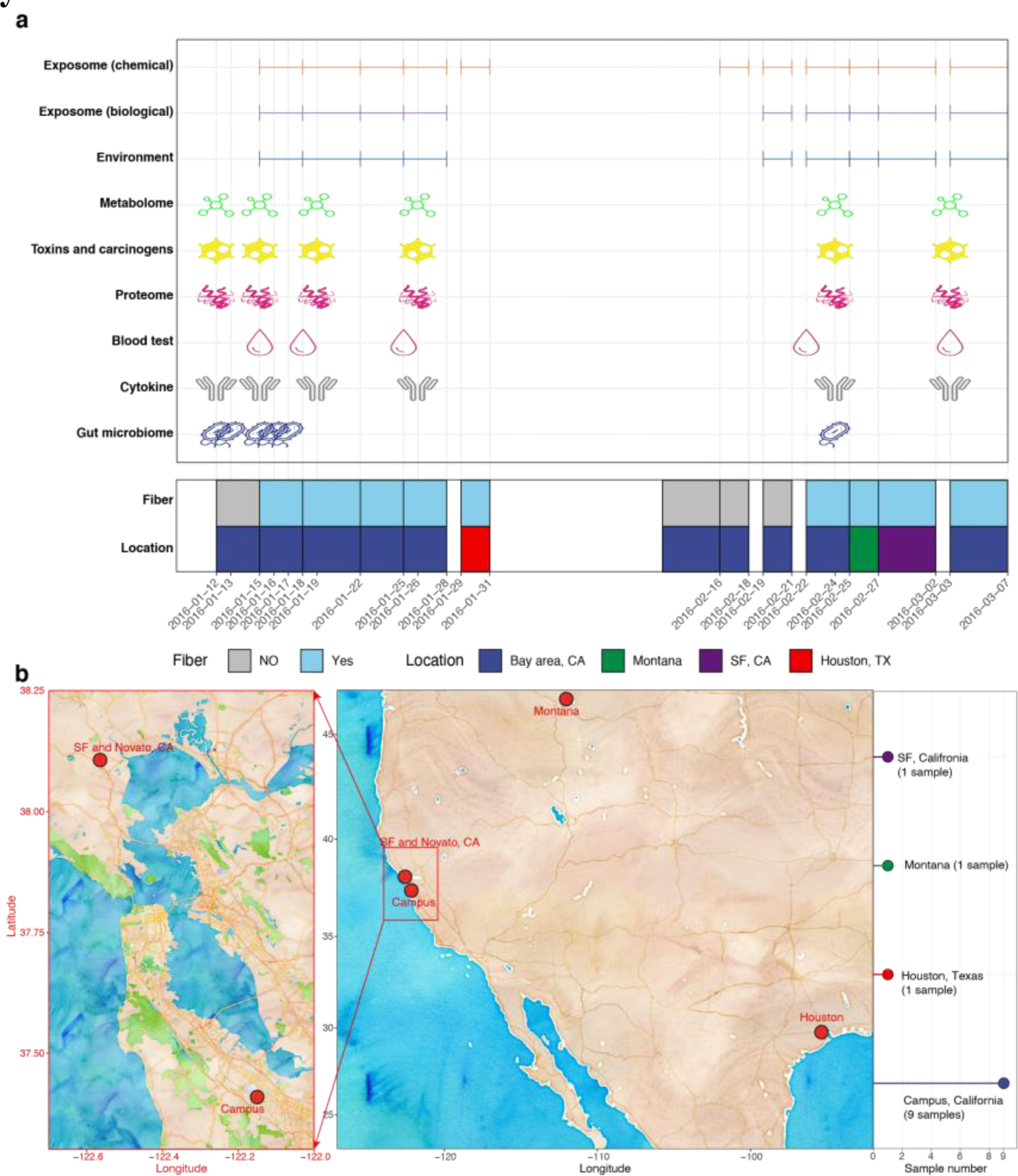
A detailed overview of sample collection. (a) Sample collection time points/periods for all exposome and internal multi-omics datasets. Corresponding fiber intake and geographical locations were also provided. (b) Collection locations for the exposome samples.

**Figure S2.**
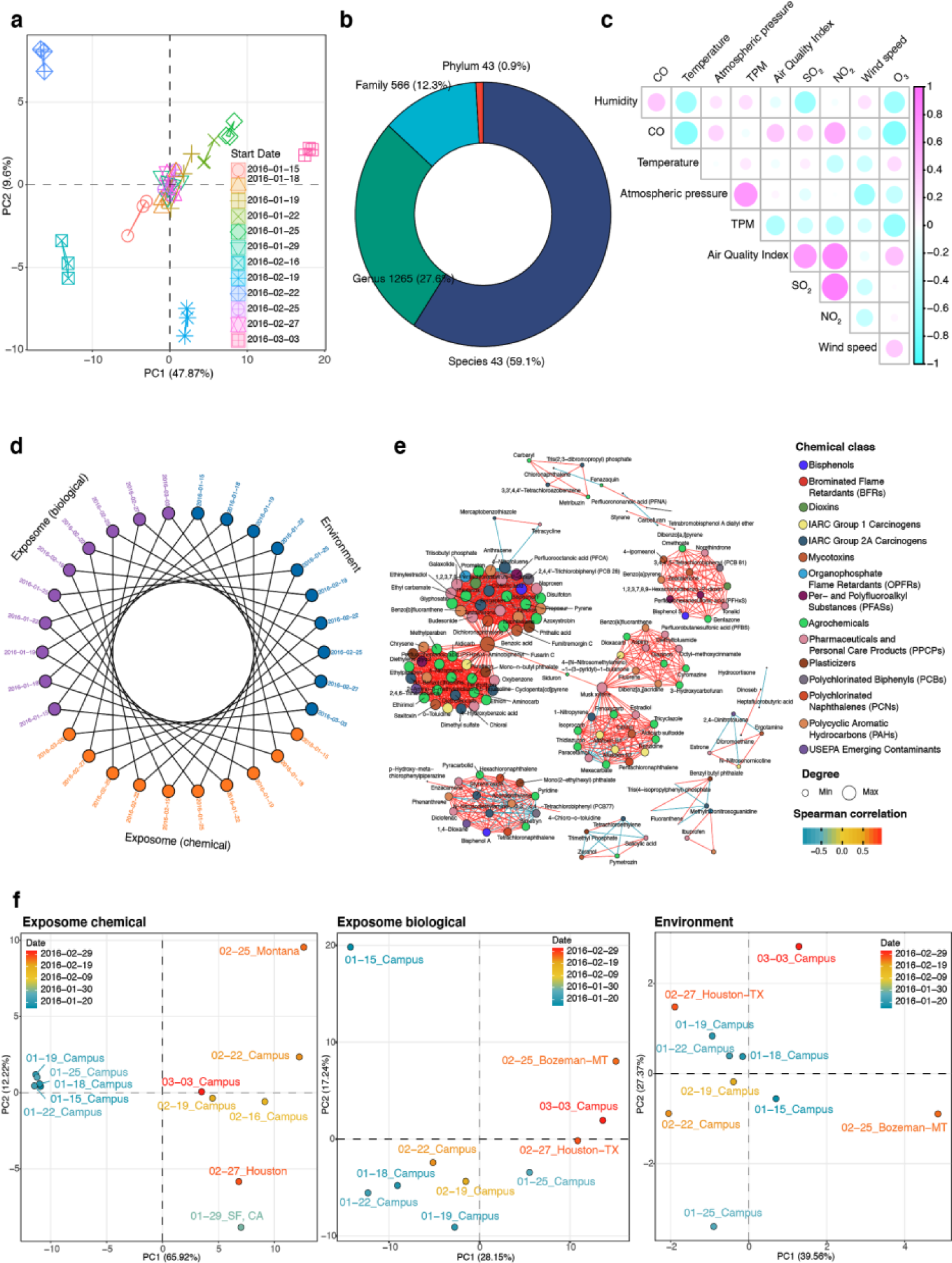
Overview of the exposome data. (a) PCA plot shows the data quality of chemical exposome data. For each sample, 3 repeats were acquired. (b) The number and percentage of biological exposome data at different taxonomic ranks. (c) Correlation plots between all environmental factors. (d) Sample matching for 3 exposome domains. (e) Intra correlation network for the chemical exposome. (f) PCA plots of the 3 exposome domains based on the sampling time and locations.

**Figure S3.**
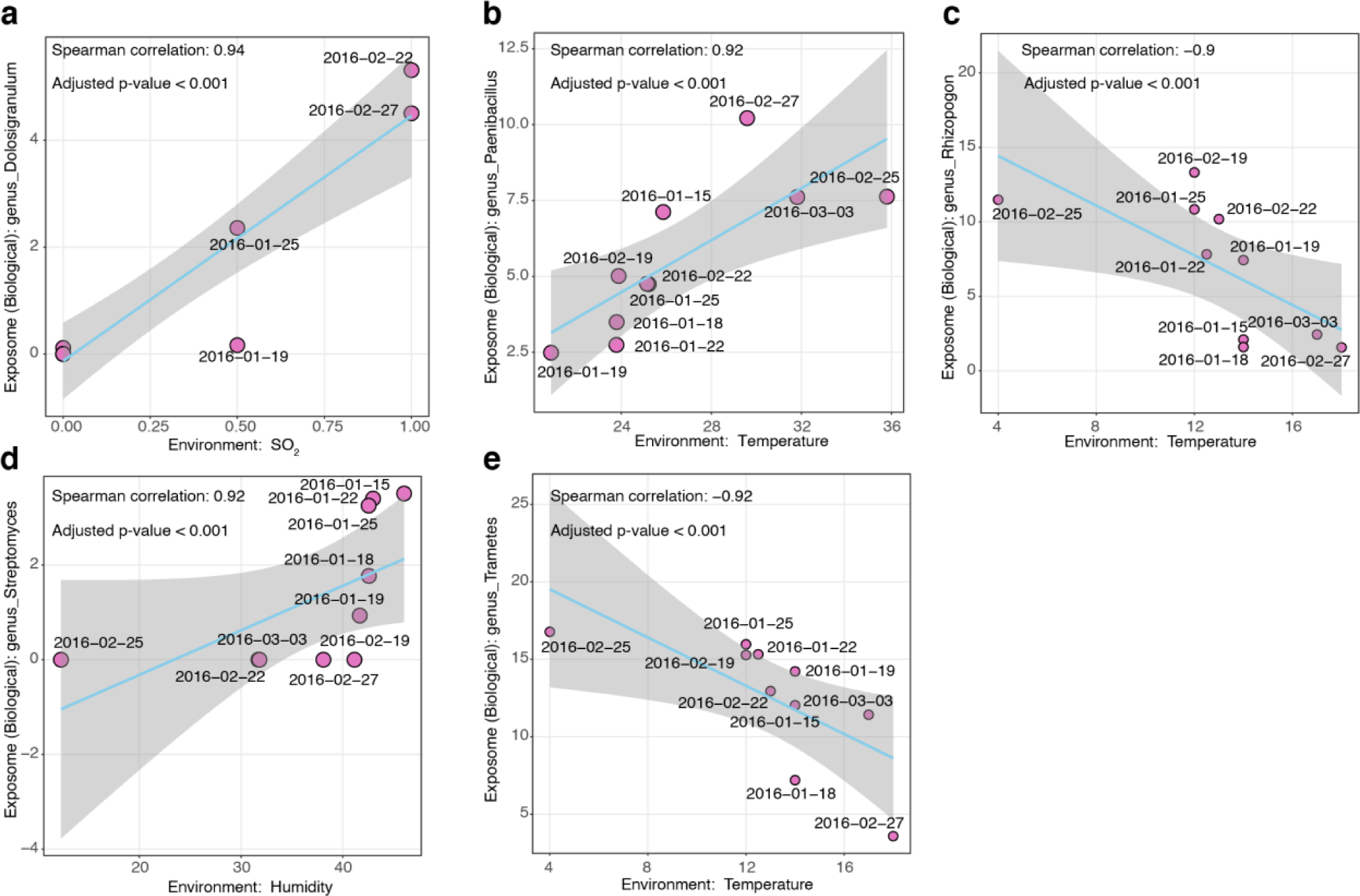
Representative Spearman correlation plots between environment factors and microbes.

**Figure S4.**
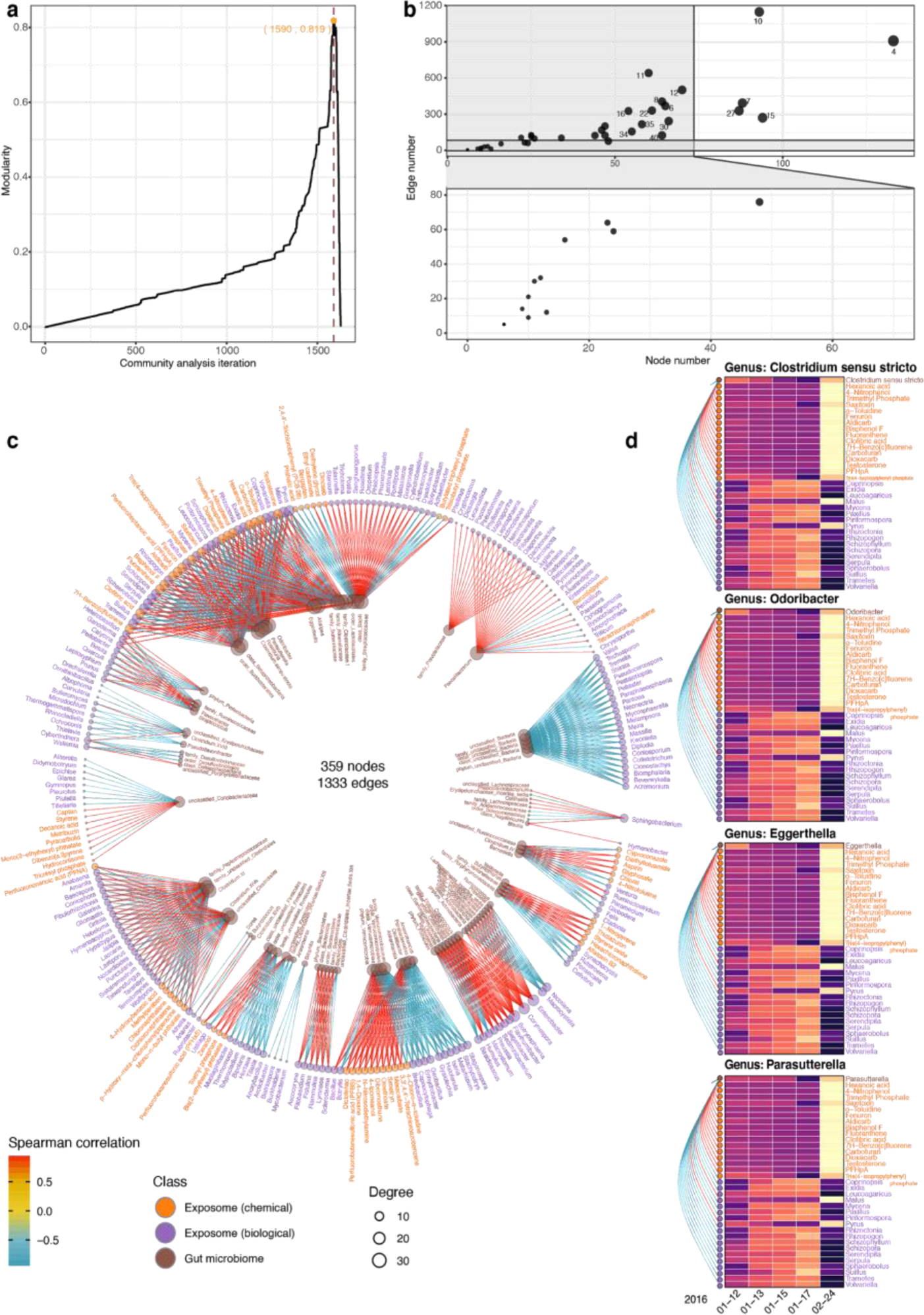
The exposome and internal multi-omics analyses. (a) The maximum modularity in the exposome and internal-omes correlation network community analysis was 0.819. (b) Node and edge numbers for all the subnetworks in the community analysis. (c) The complete correlation network between the exposome and the gut microbiome (|r| > 0.9; q-value < 0.05). (d) Representative personal gut bacteria that has 34 significant correlations with the exposome components (|r| > 0.9; q-value < 0.05).

**Figure S5.**
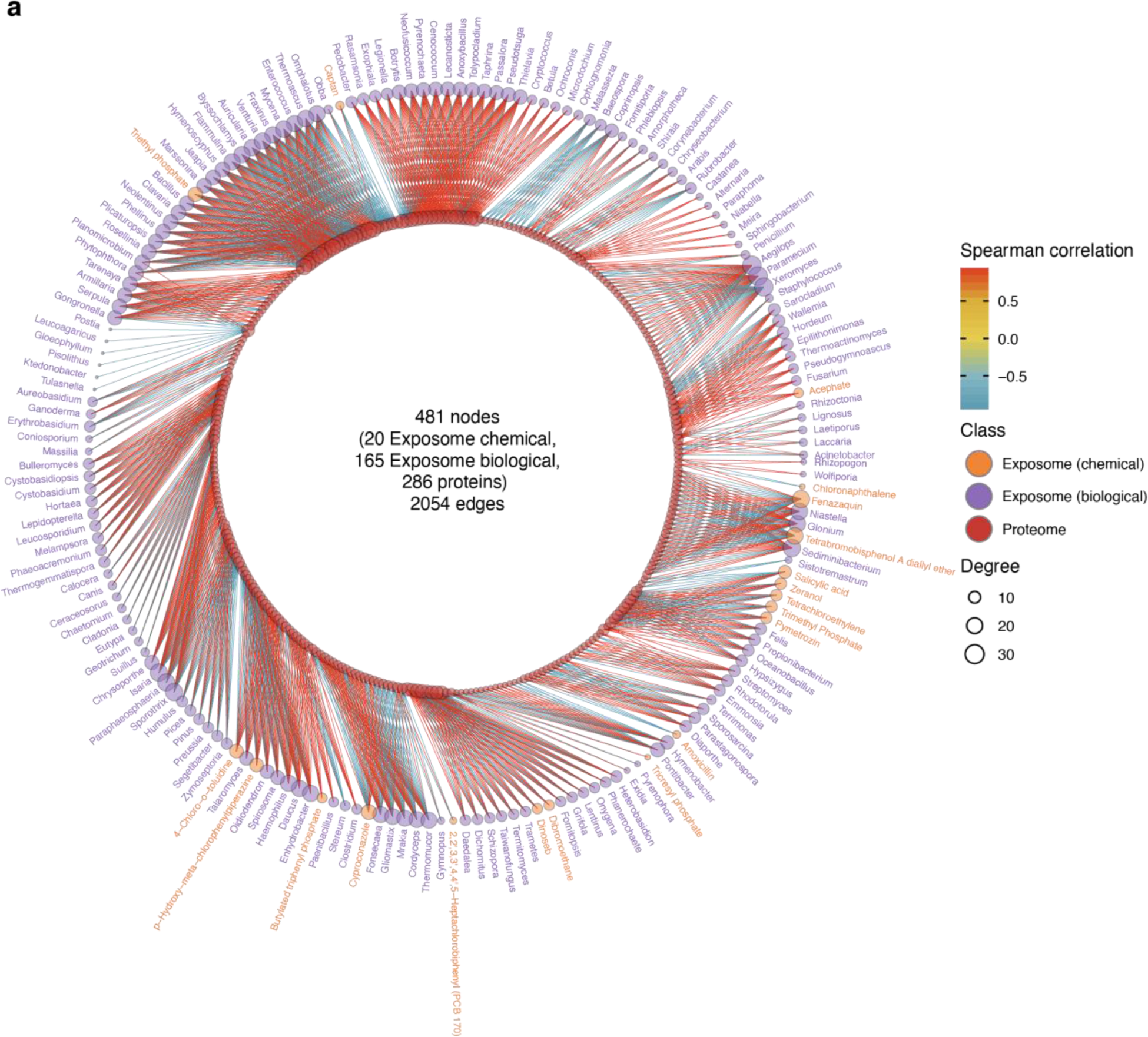
The complete correlation network between the exposome and proteome (|r| > 0.9; q- value < 0.05).

**Figure S6.**
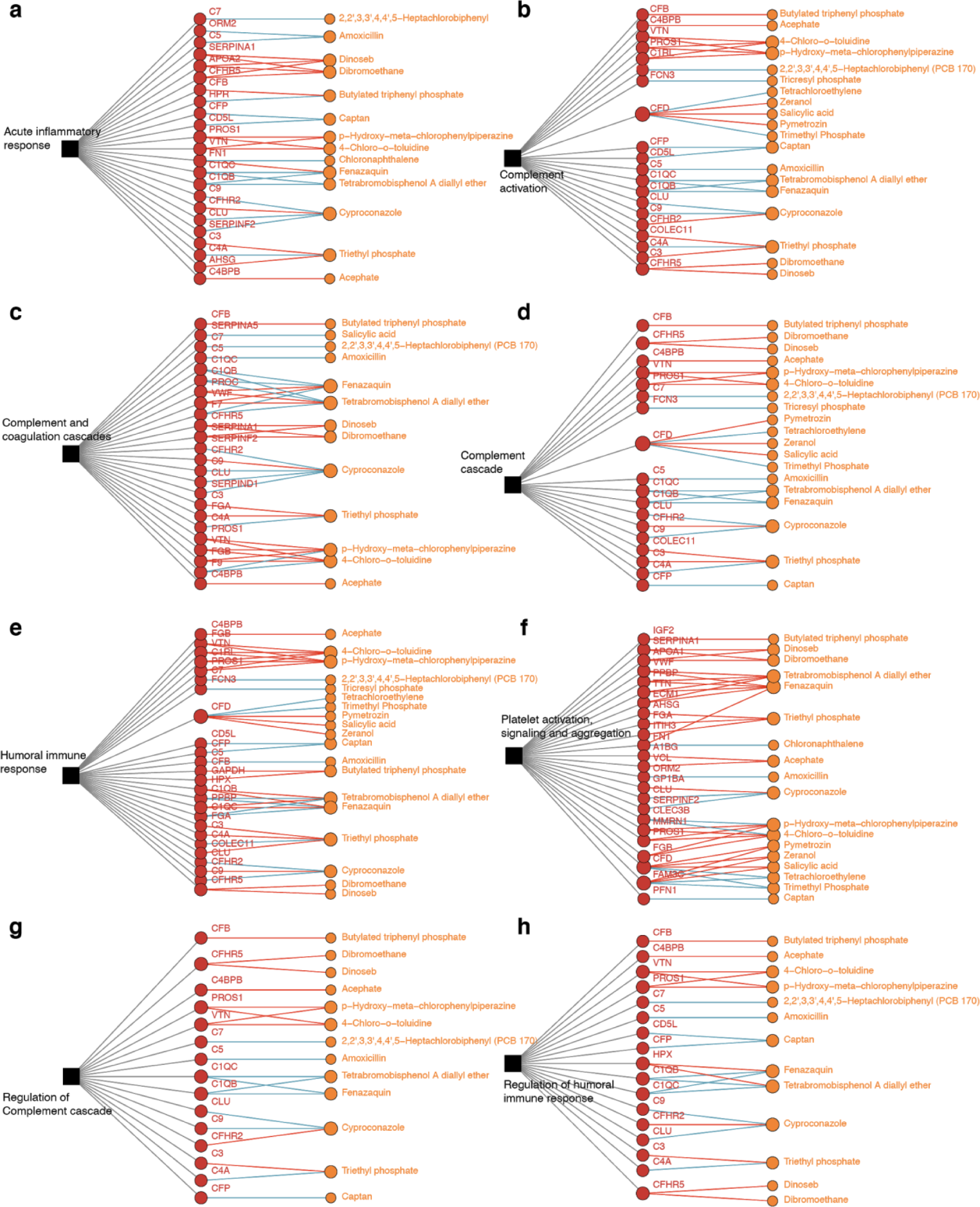
The detailed networks between the exposome and proteins for each signaling pathway (|r| > 0.9; q-value < 0.05).

**Figure S7.**
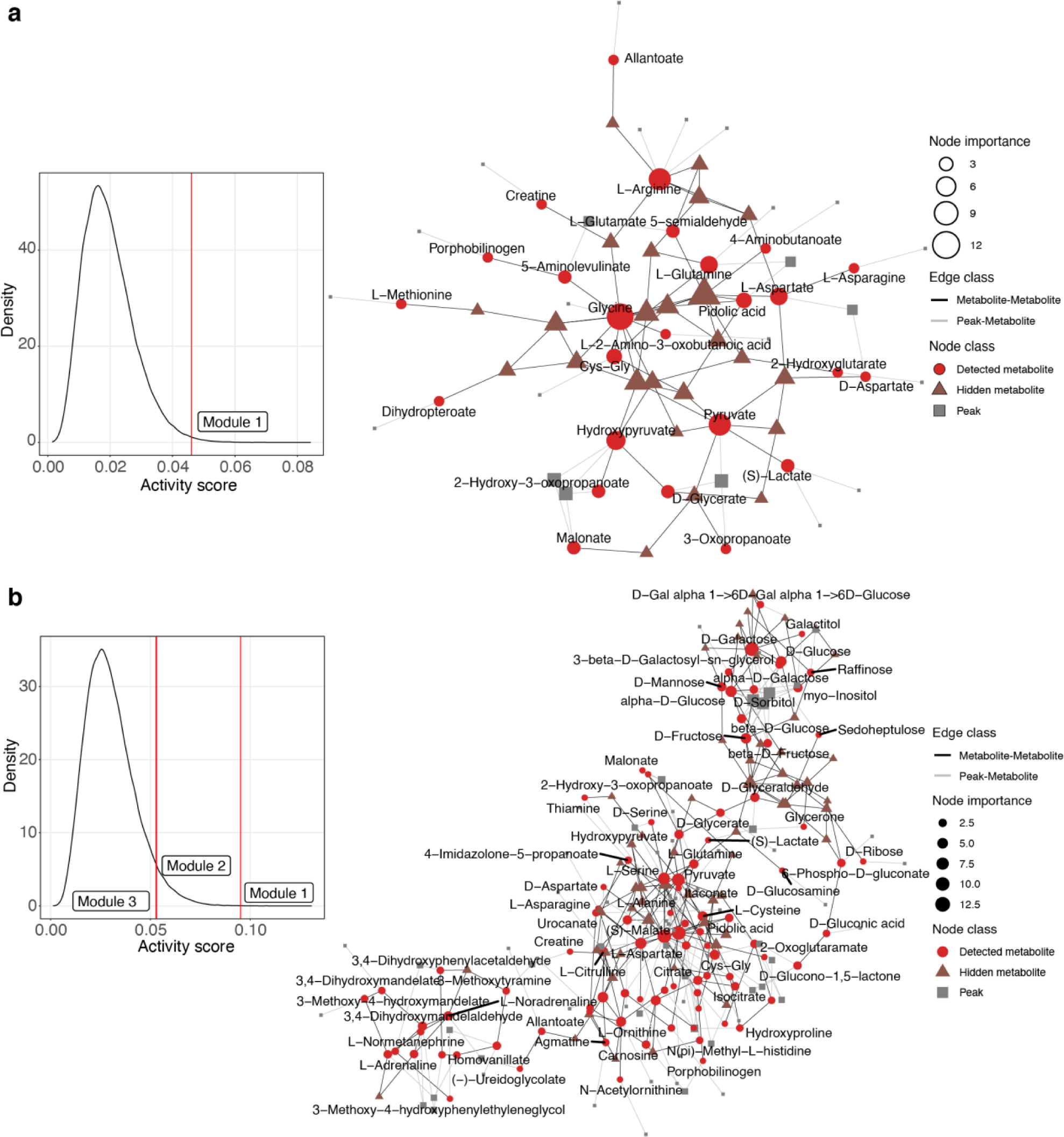
Feature-based network analysis for metabolic features significantly correlated with the exposome (**Methods**). (a) Chemical exposome. Left panel, the null distribution of activity scores and only module 1 was significant. Right panel, network constructed by significant modules. (b) Biological exposome. Left panel, the null distribution of activity scores and modules 1, 2 and 3 were significant. Right panel, network constructed by significant modules.

**Figure S8.**
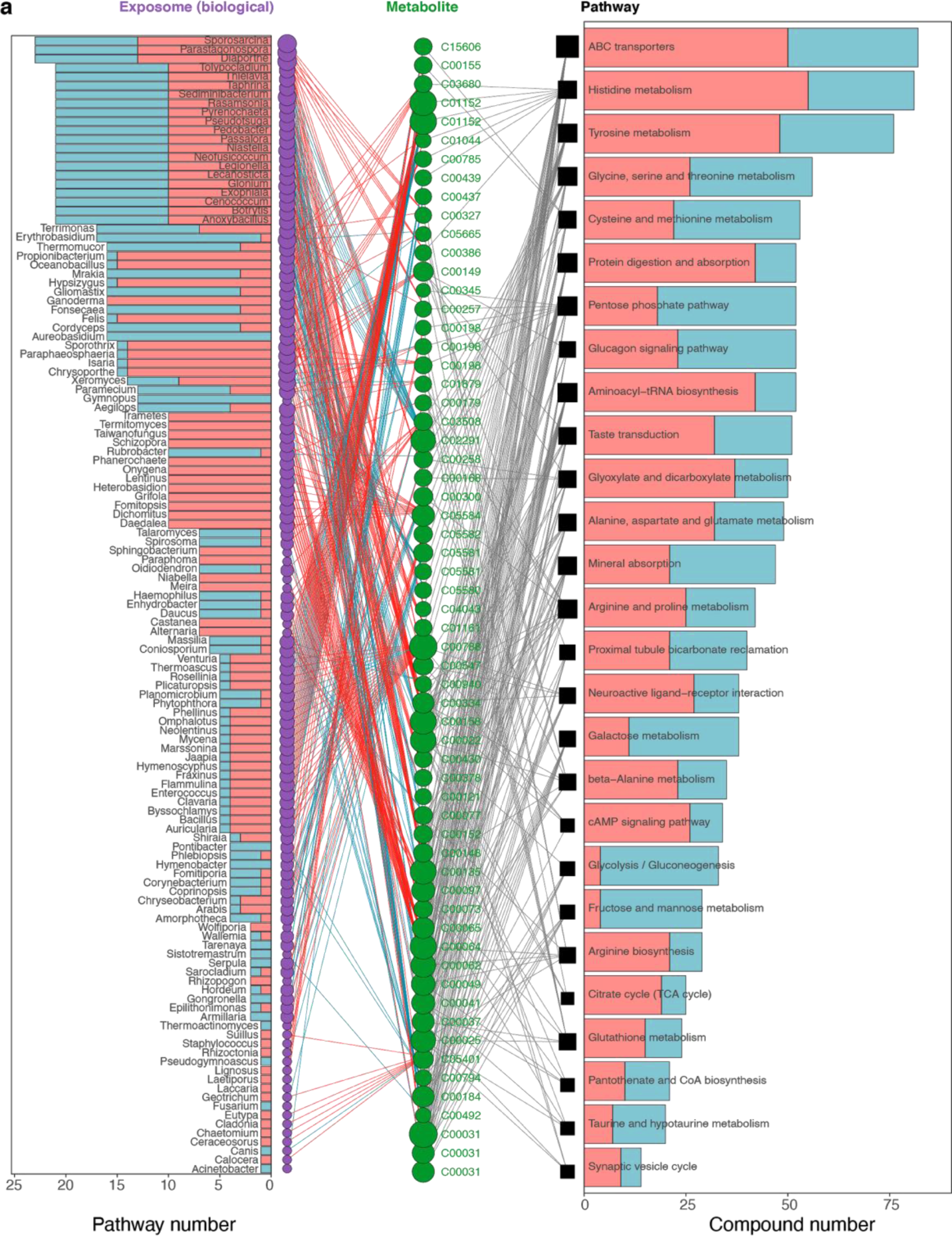
The complete network between the exposome and metabolite for each metabolic pathway (|r| > 0.9; q-value < 0.05).

## Supplementary Note

Note 1. The session information in the study

**Supplementary Table 1.**
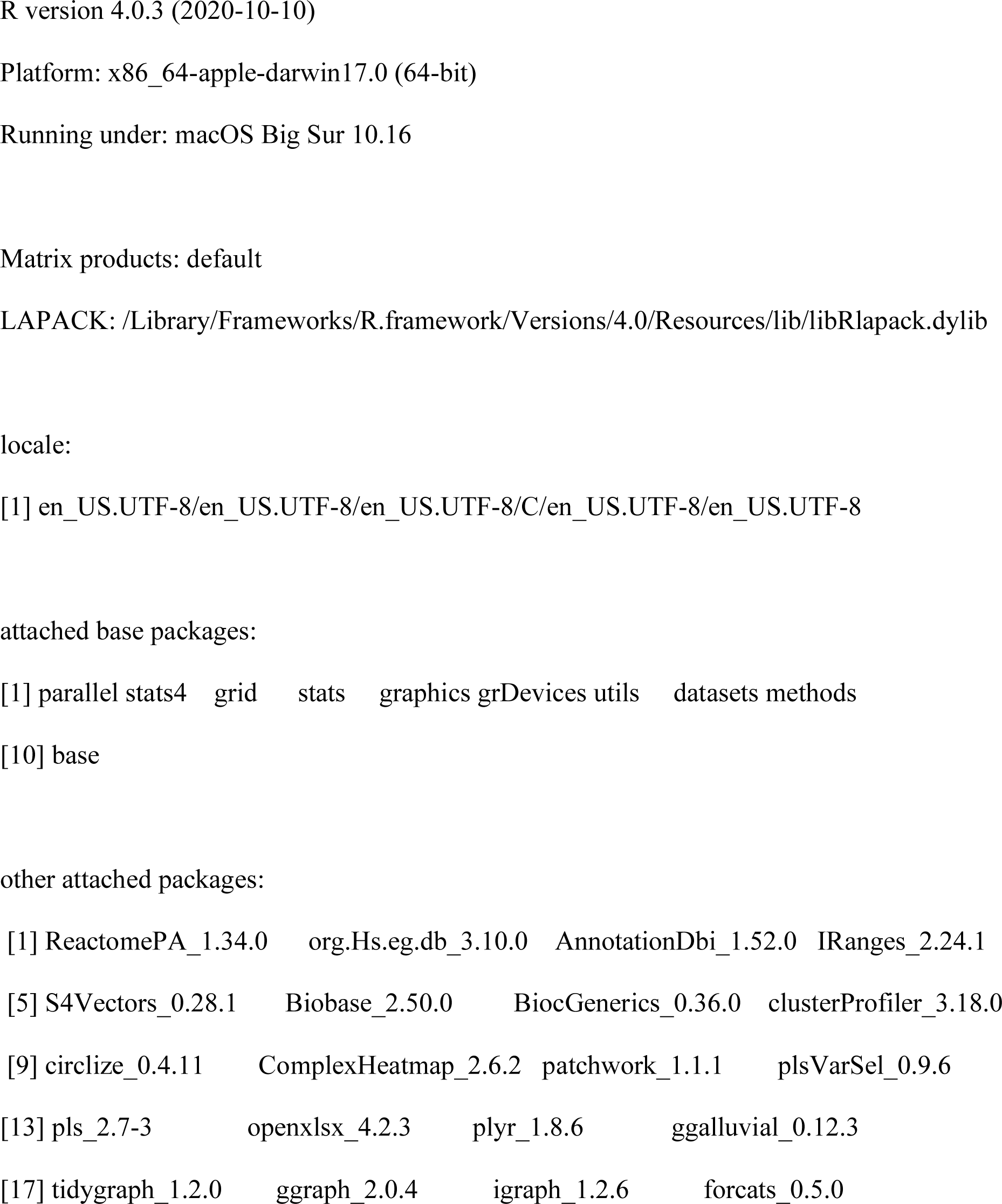
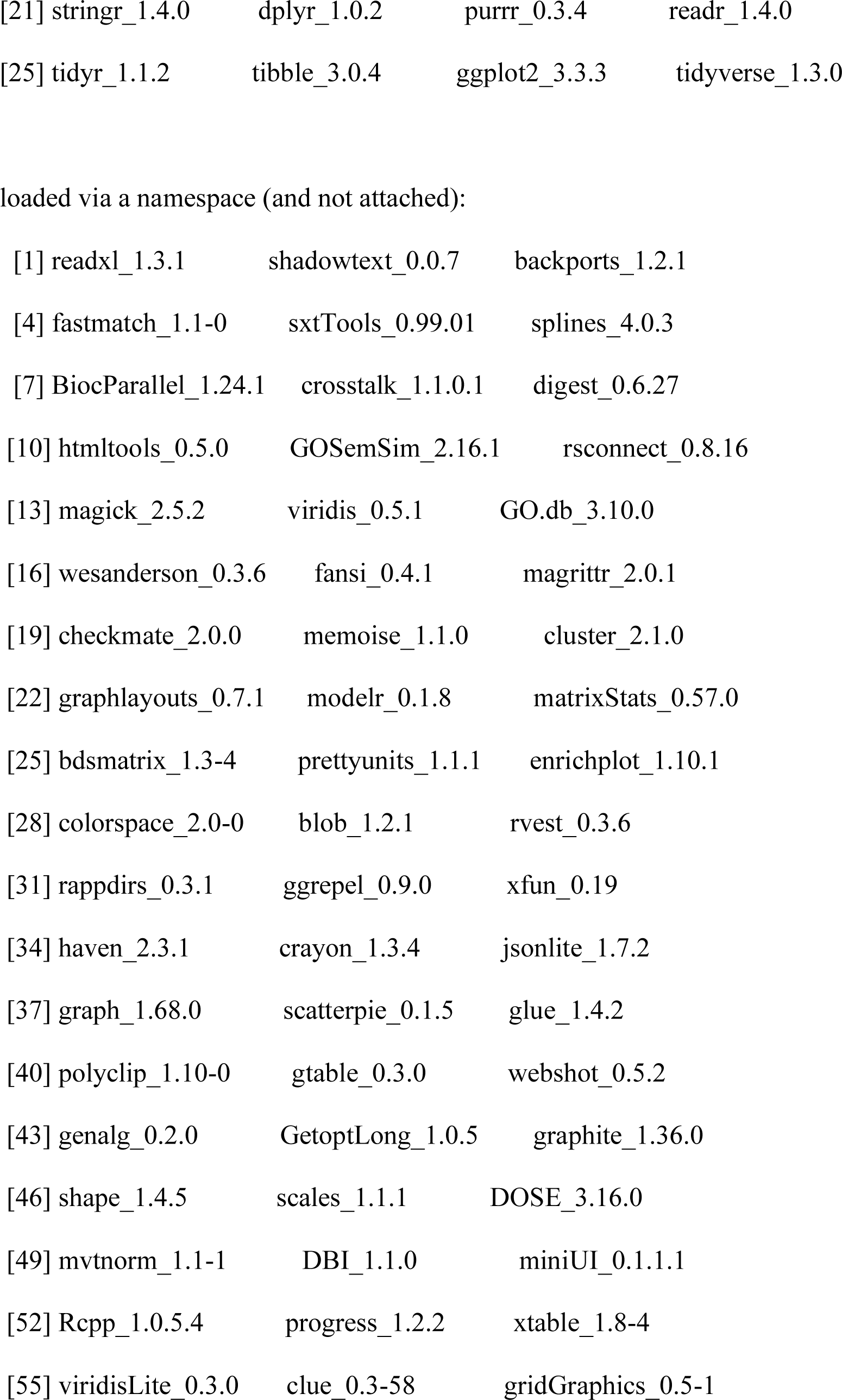
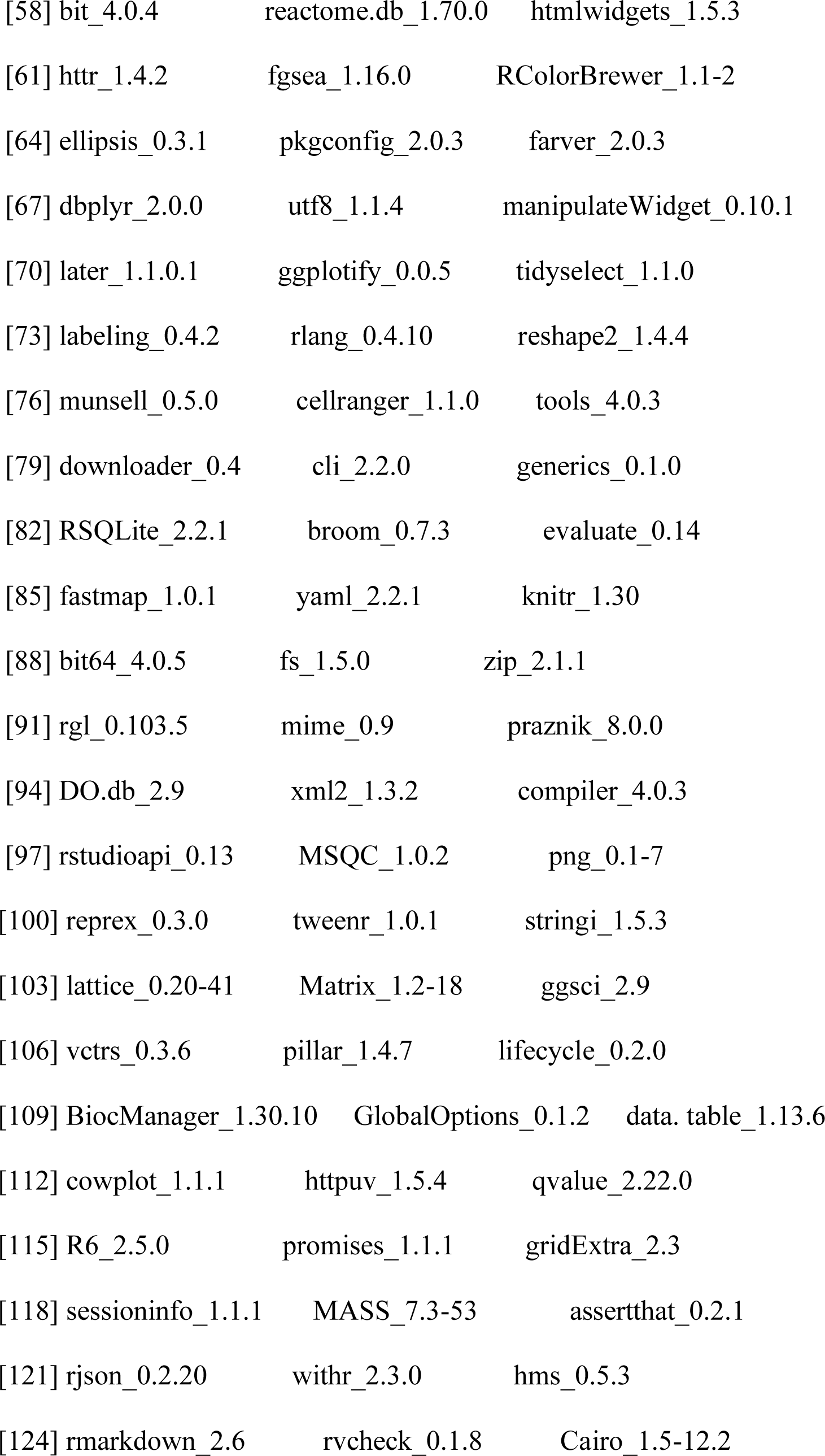
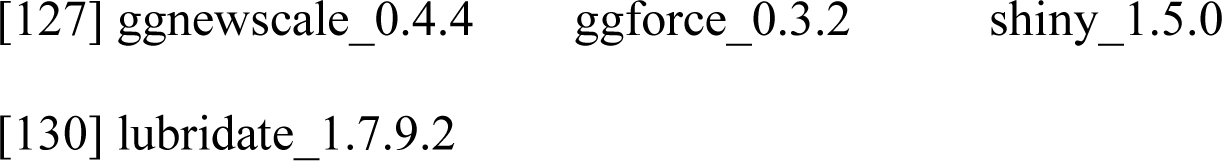

Note 2. The food log (fiber intake) of the participant in this study

From 1/15/2016 to 1/31/2016, the participant took 20 grams of arabinoxylan daily. From 2/22/2016 to 3/17/2016, the participant took 10 grams of guar gum daily.

**Supplementary Table 1.** Adduct list for metabolite annotation in this study.

| Mode | Adduct |
| --- | --- |
| RPLC positive mode | $(M+H)^+$, $(M+H-H_2O)^+$, $(M+H-2H_2O)^+$,<br>$(M+NH_4)^+$, $(M+Na)^+$, $(M-H+2Na)^+$, $(M-2H+3Na)^+$, $(M+K)^+$, $(M-H+2K)^+$, $(M-2H+3K)^+$, $(M+CH_3CN+H)^+$,<br>$(M+CH_3CN+Na)^+$, $(2M+H)^+$, $(2M+NH_4)^+$,<br>$(2M+Na)^+$, $(2M+K)^+$, $(M+HCOO+2H)^+$ |
| RPLC negative mode | $(M-H)^-$, $(M-H_2O-H)^-$, $(M+Na-2H)^-$, $(M+K-2H)^-$, $(M+NH_4-2H)^-$, $(2M-H)^-$, $(M+F)^-$ |
| HILIC positive mode | $(M+H)^+$, $(M+H-H_2O)^+$, $(M+H-2H_2O)^+$,<br>$(M+NH_4)^+$, $(M+Na)^+$, $(M-H+2Na)^+$, $(M-2H+3Na)^+$, $(M+K)^+$, $(M-H+2K)^+$, $(M-2H+3K)^+$, $(M+CH_3CN+H)^+$, |
| | $(M+CH_3CN+Na)^+$, $(2M+H)^+$, $(2M+NH_4)^+$,<br>$(2M+Na)^+$, $(2M+K)^+$, $(M+HCOO+2H)^+$ |
| HILIC negative mode | $(M-H)^-$, $(M-H_2O-H)^-$, $(M+Na-2H)^-$, $(M+K-2H)^-$,<br>$(M+NH_4-2H)^-$, $(2M-H)^-$, $(M+CH_3COO)^-$ |

## Supplementary Data

Supplementary Data 1. All exposome and internal multi-omics data used in this study.

Supplementary Data 2. The detailed information of the intra-correlation network of the chemical exposome.

Supplementary Data 3. The detailed information of the personal exposome cloud.

Supplementary Data 4. The detailed information of the inter-correlation network of the exposome and internal-multi-omics data.

Supplementary Data 5. Pathway enrichment results for proteins connected with the chemical exposome.

Supplementary Data 6. Pathway enrichment results for proteins connected with the biological exposome.

Supplementary Data 7. Pathway enrichment results for metabolic peaks connected with the chemical exposome.

Supplementary Data 8. Pathway enrichment results for metabolic peaks connected with the biological exposome.

Supplementary Data 9. The comparison between metabolite annotations from traditional methods and metabolic feature-based network analysis.

Supplementary Data 10. The contributions of the exposome on blood tests and cytokines.

## Notes

### Competing Interest Statement

The authors have declared no competing interest.

